# Transcriptomic characteristics along the longitudinal axis of the hippocampus and medial entorhinal cortex across species

**DOI:** 10.64898/2026.08.26.746438

**Authors:** Sihui Cheng, Qirun Wang, Ye Feng, Cong Chen, Guge Suri, Yilan Liang, Yikai Yang, Kai Gao, Menno P. Witter, Huixia Yang, Jie Lin, Chenglin Miao

**Affiliations:** Peking-Tsinghua Center for Life Sciences, Peking University; Beijing, 100871, China; Center for Quantitative Biology, Peking University; Beijing, 100871, China; Department of Obstetrics Gynecology and Reproductive Medicine, Peking University First Hospital; Beijing, 100871, China; School of Life Sciences, Peking University; Beijing, 100871, China; Peking-Tsinghua-National Institute Biological Sciences, Peking University; Beijing, 100871, China; The IDG/McGovern Institute for Brain Research, Peking University; Beijing, 100871, China.; Kavli Institute for Systems Neuroscience, Norwegian University of Science and Technology, Trondheim, 7030, Norway

## Abstract

The hippocampus (HPC) and medial entorhinal cortex (MEC) are essential for learning, memory, and spatial cognition, and both exhibit dorsoventral (longitudinal) organization across mammalian species. While prior studies have highlighted functional differences along this axis, the molecular basis and cross-species conservation of these differences remain poorly understood. Here, we employed spatial transcriptomics to generate a comprehensive molecular atlas of HPC and MEC in five species—human, tree shrew, mouse, canine, and pig—standardizing the dorsoventral axis for cross-species comparison. Using support vector machine (SVM) models, we identified high-weight genes predictive of dorsoventral identity and revealed conserved functional patterns: dorsal HPC was enriched for cytoskeletal and synaptic pathways, while ventral HPC favored nucleotide and energy metabolism. In the MEC, dorsal regions were enriched for calcium transport and lipid metabolism, whereas ventral regions were associated with calcium homeostasis and amyloid regulation. Species-specific SVM models uncovered dramatic divergence, leading us to propose the ancestral confinement theory, suggesting that conserved dorsoventral features are maintained within an evolutionary framework that permits species-specific adaptations. To link molecular patterns with cell types, we conducted single-nucleus RNA sequencing of tree shrew HPC and MEC and integrated data from other species. Deconvolution analysis showed species-specific GABAergic neuron distributions along the axis, with notable dorsal enrichment in human and ventral enrichment in other species. Together, our findings provide a cross-species molecular framework of HPC and MEC organization, revealing both conserved and species-specific dorsoventral programs underlying brain function and evolution.

## Introduction

The HPC and MEC are key brain regions involved in cognition^1^ . They show remarkably conserved structures and functions during evolution across mammalian species^2^. A fundamental organizational principle of these structures is their functional segregation along the dorsoventral (septotemporal) axis^3^. In rodents, the dorsal HPC is crucial for spatial tasks, as its lesions impair water maze performance^4^, while ventral HPC lesions do not^5,6^. The dorsal HPC shows a higher density of place fields. Conversely, the ventral HPC is more associated with emotional regulation. Lesions in the ventral HPC increase stress responses and decrease anxiety, as shown in elevated plus maze and stress studies^5,6^. The ventral HPC also plays a role in fear conditioning^6^, affecting tone fear more than context fear, likely due to its direct connections with the amygdala^2^. A similar functional topography exists along the longitudinal (anteroposterior) axis of the human HPC, with the anterior part corresponding to the ventral HPC and the posterior part corresponding to the dorsal HPC in rodents. Inactivation of the right dorsal HPC disrupts spatial memory retrieval in rodents, consistent with findings in human where the right posterior HPC is activated during route recall^7–10^.

This functional segregation is paralleled by anatomical connectivity patterns. In rodents, the dorsal CA1 region of the HPC, rich in place cells, projects to the subiculum and related areas. This network is key for navigation and memory, connecting to the retrosplenial and anterior cingulate cortices. It also links to subcortical structures, such as the mammillary bodies, crucial for spatial mapping and exploratory behavior^11,12^. The ventral HPC connects with the olfactory bulb and amygdala, influencing emotional regulation and neuroendocrine functions^2^. It projects to the hypothalamus, governs behaviors including feeding and defense, and connects with the nucleus accumbens and circadian rhythm structures, affecting reward, motivation, and sleep^11,12^. The human HPC is not functionally uniform along its longitudinal (anteroposterior) axis either. A compelling body of evidence from structural and functional neuroimaging^13^, as well as studies of patients with lesions^14^, supports a functional dissociation: the posterior HPC is preferentially involved in fine-grained, high-resolution spatial and contextual representations, enabling precise navigation and detailed memory. In contrast, the anterior HPC is implicated in broader, coarser representations, supporting gist-based memory, context generalization, and emotional valence processing. This gradient is thought to reflect differential connectivity patterns^15^, with the posterior HPC more connected to cortical regions involved in visual-spatial processing (e.g., retrosplenial cortex, posterior parietal cortex) and the anterior HPC more connected to affective and interoceptive regions (e.g., amygdala, anterior cingulate, orbitofrontal cortex).

The entorhinal cortex (EC) has a dorsal ventral axis as well^2^, which can be divided into three parallel zones: caudolateral, intermediate, and rostromedial. The caudolateral zone receives visuospatial inputs, projecting to the dorsal HPC. The medial zone, with olfactory, visceral, and gustatory inputs, connects to the ventral HPC. The intermediate zone, with widespread inputs, projects to intermediate hippocampal regions. These projections follow a topographic pattern throughout the trisynaptic circuit, conserved across species. The EC innervates all hippocampal components; CA1 and the subiculum send reciprocal projections back to the EC, maintaining the same topographic organization^2^. The MEC is related to spatial memory with a high percentage of grid cells^1^, likely associated to its specific inputs from the presubiculum, where one finds a predominance of head direction cells^16,17^ as well as other spatially modulated neurons^18^. Consistent with the connection of MEC to HPC, it also has a functional diversity along the dorsoventral axis. The grid cells, believed to be the major component of the spatial map, exhibit a spatial scale along the dorsal-ventral axis^19^.

This functional topography is thought to be supported by graded gene expression patterns^20,21^, yet these molecular architectures in MEC and the extent to which they are conserved across phylogenetically diverse species remains poorly understood. Animal studies, particularly in rodents, have explored the cellular and molecular segregation in the HPC along the dorsal to ventral axis^22–26^, but similar studies in the MEC are still lacking. In addition, most of these studies did not systematically investigate the cross-species difference and conservation of expression profiles in HPC and MEC. It would be essential to study the molecular and cellular changes of HPC and MEC across mammalian species during evolution. Furthermore, previous studies also have limitations with their techniques. For example, the large-scale in situ hybridization (ISH) identified gene expression differences along the mouse dorsal-to-ventral axis^23,24^, but with a limited number of genes analyzed. The bulk RNA sequencing^25^ and single-nuclei RNA-sequencing (snRNA-seq)^26^ have identified major cell-class transcriptomes but lacked spatial resolution. The recently developed spatial transcriptomics technologies represent significant methodological advancements, providing both larger-scale profiling capabilities and substantially improved spatial resolution compared to conventional approaches^27–29^.

In this study, we investigate cell-type-specific variability of expression profiles in 5 species along the dorsal to ventral axis of HPC and MEC, using 10X Spatial transcriptomics and 10X HD Spatial transcriptomics, with high spatial resolutions both at the anatomical and cellular levels. This study included human fetuses at 32 weeks of gestational age as research subjects, a developmental stage at which the HPC and MEC are relatively well developed in longitudinal axis^30,31^, and preserved gene expression patterns along this axis becoming increasingly stable^32–34^. The tree shrew was also included as a study species, which resembles a squirrel in appearance, exhibits remarkable similarities to primates—and even human—in terms of neurodevelopment, neuroanatomy, and stress-related behavioral responses. Moreover, tree shrew possess the highest brain-to-body weight ratio among mammals, making them one of the ideal models for investigating brain function^35^. Given that most previous research on dorsoventral axis connectivity and functionality of the HPC and MEC has primarily relied on mouse models, mice were also included here to enable effective cross-species comparisons. Beagle canine, internationally recognized as standard laboratory canine, were selected as they share similar capacities for emotional and social information processing, making them an increasingly important model in neuroscience research^36^. Finally, domestic pig of the Duroc × Landrace × Yorkshire (DLY) hybrid breed was included. Compared to smaller animal models, pig possess a brain that is more similar to the human brain in both size and morphology, exhibiting a complex gyrencephalic structure, thus providing another valuable species for cross-species comparative analyses^37^. Together, we selected human, tree shrew, mouse, canine, and pig to capture both phylogenetic diversity and translational relevance. This combination spans key mammalian lineages, from rodents and primates to large-brained gyrencephalic species, enabling the identification of conserved dorsoventral programs and species-specific adaptations.

## Results

### Spatial transcriptomics of HPC and MEC from five species

We profiled the spatial transcriptomics of fresh-frozen samples from the tree shrew (*Tupaia chinensis*), mouse (*Mus musculus*), pig (*Sus scrofa*), and canine (*Canis lupus familiaris*) using 10X Visium, as well as fresh-paraffin-embedded samples from a 32-week human fetus (*Homo sapiens*) with 10X Visium HD (Figure 1A, Figure S1, S2, and S3). After quality control (Method), a total of 65,749 spots were obtained. Subregions, including CA1 (Cornu Ammonis 1), CA2 (Cornu Ammonis 2), CA3 (Cornu Ammonis 3), Or (Stratum Oriens), Rad (Stratum Radiatum), and LMol (Stratum Moleculare) from the cornu ammonis area of the HPC; PoDG (Polymorphic Layer of the Dentate Gyrus), GrDG (Granule Cell Layer of the Dentate Gyrus), and Mol (Molecular Layer) from the dentate gyrus of the HPC; and MECde (Medial Entorhinal Cortex, Deep Layers, including LIV, LV, and LVI), MECint (Medial Entorhinal Cortex, Intermediate Layers, including LII and LIII), and MECsu (Medial Entorhinal Cortex, Superficial Layers, including LI) from the MEC were manually registered based on anatomical structures (Figure 1A, Figure S1, S2, S3, and Method). Notably, we did not distinguish the CA4 (Cornu Ammonis 4) region, as studies suggest CA4 is not distinct from PoDG in terms of cellular composition^38,39^. We identified 13,726 one-to-one orthologous genes across the five species, which were used in subsequent analyses (Figure S4A and B).

**Figure 1:**
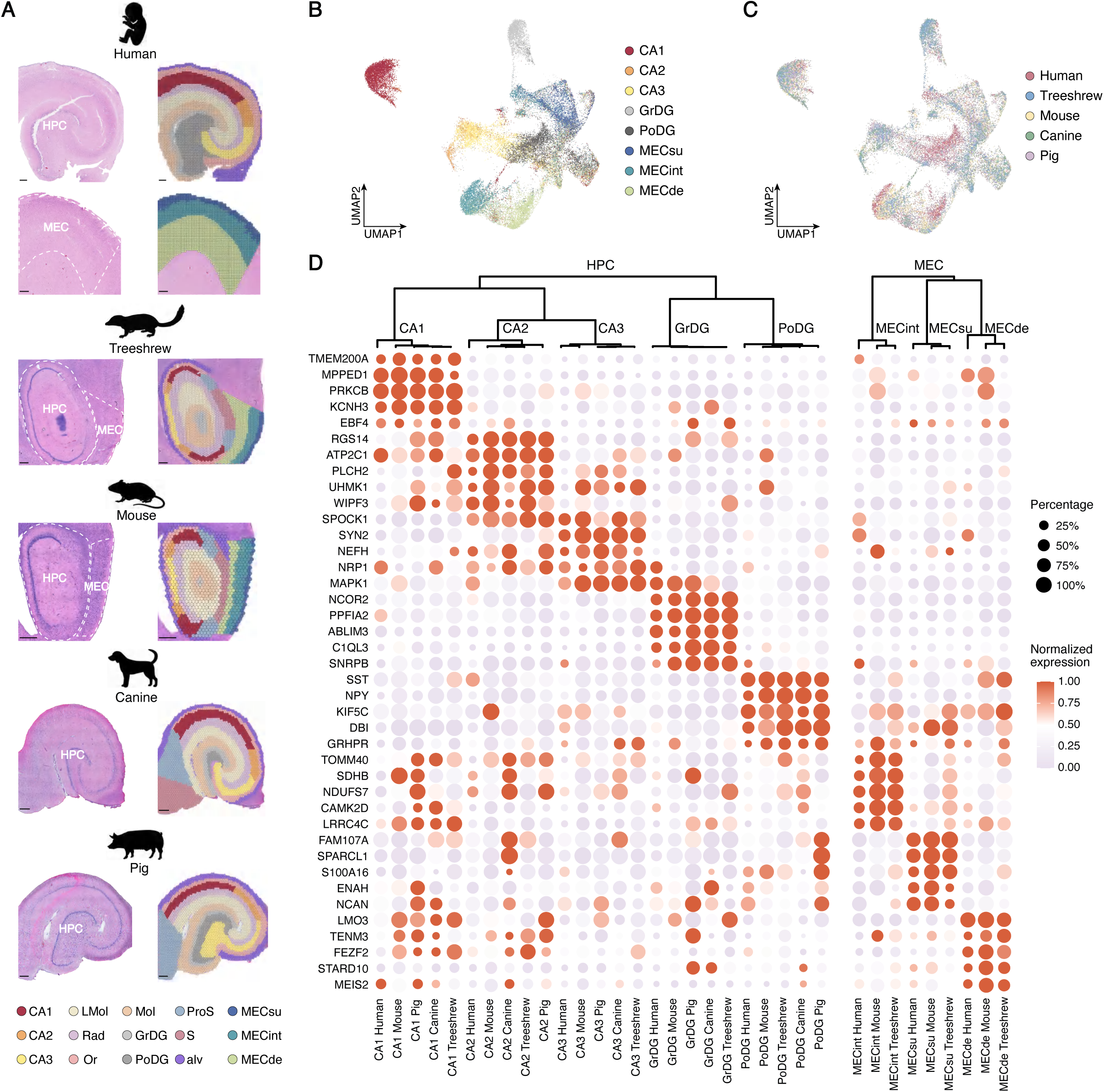
Spatial transcriptomics of HPC and MEC from five species. (**A**) Schematic representation of the samples and anatomical subregions of the HPC and MEC. (**B**) Uniform Manifold Approximation and Projection (UMAP) visualization of the integrated data from neuron-rich subregions across five species, with subregions labeled. (**C**) UMAP visualization as in (B), but with species labeled. (**D**) Gene expression profiles of the integrated data from neuron-rich subregions. The upper panel shows hierarchical clustering of the data grouped by species and subregion. The lower panel highlights marker genes for each group.

To gain a comprehensive overview, we integrated data from neuron-rich subregions of the HPC (CA1, CA2, CA3, PoDG, and GrDG from all 5 species) and the MEC (MECde, MECint, and MECsu from 3 species) using Seurat integration^40^ across species and visualized the results with uniform manifold approximation and projection (UMAP). Subregions tended to cluster together (Figure 1B), while species were uniformly distributed (Figure 1C), indicating conserved gene expression profiles within the same subregions across different species. To validate this, we performed hierarchical clustering of data grouped by species and subregions. As expected, subregions from different species clustered together (Figure 1D).

The higher resolution of Visium HD enabled finer segmentation of anatomical subregions. We examined whether the anatomically distinct layers of the MEC, including LI, LII, LIII, LV, and LVI, also exhibited transcriptomic differences. UMAP revealed distinct clustering of different layers, with specific markers identified for each layer, indicating that the layers are transcriptomically distinct (Figure S4C, D, E, and F). In addition, the data showed that subregions exhibited conserved expression profiles across species.

### Conserved dorsoventral organizational patterns across species of both HPC and MEC

Evidence suggests overarching differences along the dorsoventral axis within the HPC and MEC in both rodents and primates^2,41,42^. To investigate whether these dorsoventral differences manifest consistently across all subregions of the HPC and MEC across species, we employed the supervised machine learning method, Support Vector Machine (SVM). SVM identifies an optimal hyperplane to separate two groups of points (Figure 2A) and is well-suited for RNA sequencing data classification, where each spot represents a point in an n-dimensional space, where n is the number of genes.

**Figure 2:**
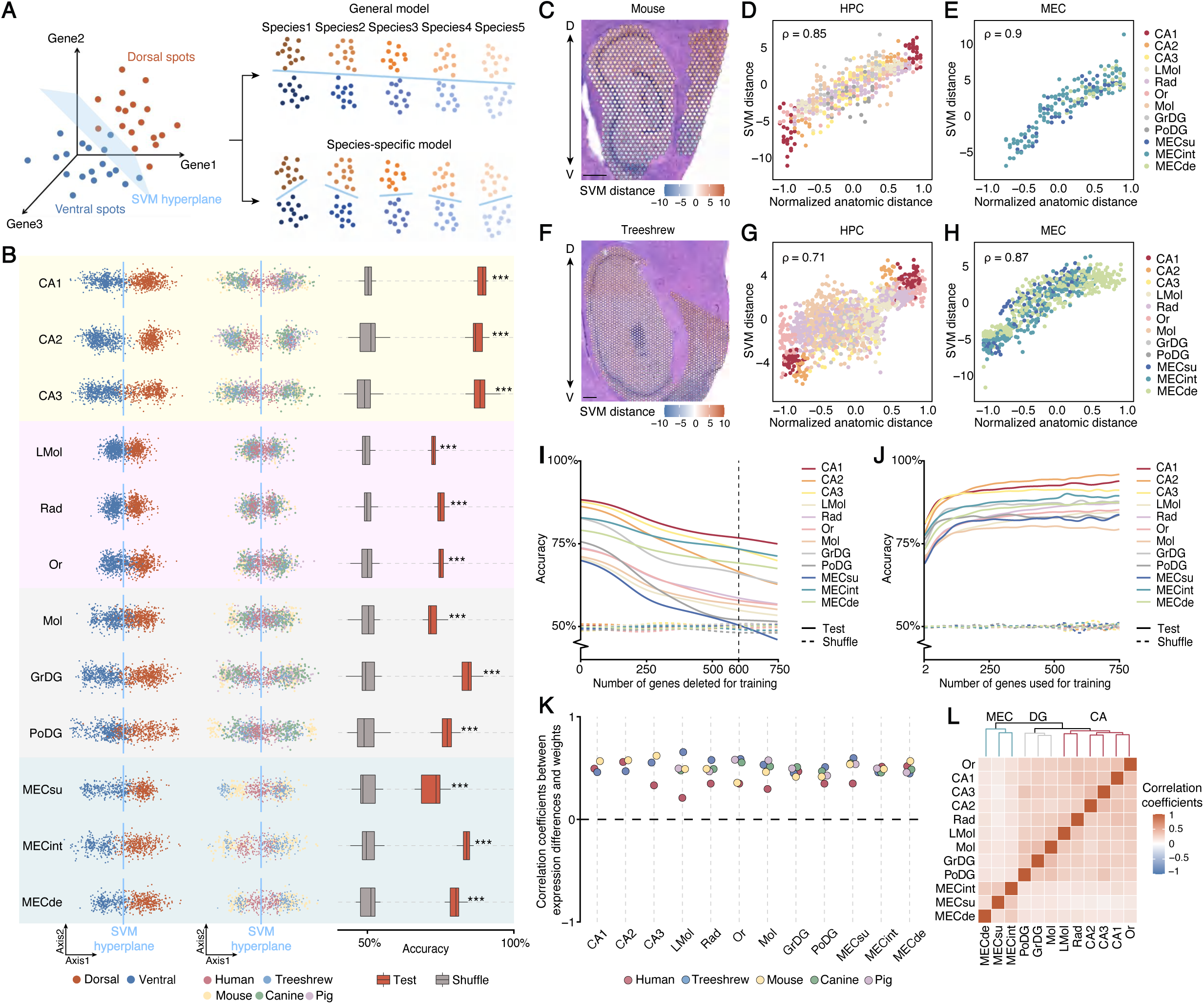
Conservative dorsoventral differences across species are evident in all subregions of HPC and MEC. (**A**) Schematic of the Support Vector Machine (SVM) methodology and training strategies for the general model and species-specific models. (**B**) Results of SVM classification using the general model. Left: dimensionality reduction visualization showing SVM-based classification of dorsoventral regions, with SVM hyperplanes indicated by light blue lines. Middle: same plot as left, labeled by species. Right: classification accuracies for each subregion. Two-sided Wilcoxon tests compare the accuracy between the testing and shuffled groups. P-values are corrected using the false discovery rate (FDR), and significance levels are indicated by asterisks, with “***” representing FDR < 0.001. (**C**) SVM distances of the general model for each spot on mouse brain sections, with correlations between normalized anatomic distances and SVM distances analyzed for each subregion of the HPC (**D**) and MEC (**E**). Pearson correlation coefficients (ρ) are calculated and tested using two-sided t-tests (p = 5.29e-152 for HPC and p = 6.87e-71 for MEC). (**F**, **G**, and **H**) Similar analysis as in (E, F, and G) conducted for tree shrew (p = 6.85e-266 for HPC and p = 2.14e-193 for MEC). (**I**) Classification accuracies of the general model when trained with the top n weighted genes removed for each subregion. (**J**) Classification accuracies of the general model when trained using only the top n weighted genes for each subregion. (**K**) Correlations between expression differences from each species and SVM weights from the general model for every subregion and species. All correlations are significantly positive (p < 0.05) as determined by two-sided t-tests. (**L**) Pairwise correlations of SVM weights from the general model across subregions.

To identify conserved dorsoventral differences across species, we aggregated data from all species and trained SVM models for each subregion. Shuffled tag data served as a control for training the models. Models trained with combined data from all species are referred to as the “general model”, while those trained with data from each species are termed “species-specific models” (Figure 2A). Dimensional reduction along the normal vector of each SVM hyperplane (Method) revealed that dorsal and ventral spots were separated by the SVM decision boundary across all species (Figure 2B, left and middle panels). Significantly higher classification accuracies are observed compared to shuffled groups (Figure 2B, right panel), confirming common dorsoventral differences among the five species. Interestingly, conserved dorsoventral differences were also observed even in white-matter-rich subregions. To validate the general models’ predictive power in other samples, we performed classification on single-nucleus RNA sequencing data from excitatory neurons of adult human^26^. Pyramidal neurons were predicted using the general models of CA1, CA2, and CA3, while DG neurons were predicted using the general models of GrDG and PoDG. Despite substantial differences in sequencing methods and sample ages, significant classification performance was achieved (Figure S5A and B), demonstrating the robustness of our models. We also validate the consistency between the model characteristics and the expression profiles of mouse HPC and MEC from multiple data sources (Figure S5C, D, and E).

To further validate the conserved dorsoventral heterogeneities in gene expression profiles and mitigate potential SVM bias, we conducted additional analyses using linear discriminant analysis (LDA), another supervised machine learning method for classification. The results showed strong correlation to the results of SVM prediction (Figure S5F) and significantly higher accuracies across all regions compared to shuffled groups (Figure S5G), confirming conserved dorsoventral heterogeneities among species.

Given the continuous variation in gene expression along the dorsoventral axis in the HPC^2^, we inferred that if SVM models capture true differences in expression profiles, the distances of spots from the hyperplane in high-dimensional space should reflect real anatomical distances in the brain. Indeed, predicted distances for each subregion correlated with anatomical positions, with greater dorsal or ventral positions corresponding to larger predicted distances (Figure 2C-H and Figure S5H).

Next, we examined which genes contribute to the conserved dorsoventral differences across species within each subregion. One advantage of SVM is its interpretability. Once the hyperplane is determined, the weights of all genes are established. The more weight the gene has, the more the gene contributes to the classification. To assess how many genes underlie the general dorsoventral differences, we trained SVM models under two conditions: (1) with the top n weighted genes removed, and (2) using only the top n weighted genes. We found that removing up to approximately 600 top-weighted genes still yielded classification accuracies above those of the shuffled control group across all subregions (Figure 2I). Conversely, using only approximately the top 100 weighted genes was sufficient to achieve high classification accuracies for all subregions (Figure 2J). These findings suggest that a few hundred top-weighted genes are essential for defining the conserved dorsoventral axis across species.

The weights assigned to genes by the SVM model are expected to reflect their expression patterns along the dorsoventral axis: positively weighted genes should be more highly expressed in dorsal regions, while negatively weighted genes should be more highly expressed in ventral regions. To validate this interpretation, we computed the correlation between gene weights and their dorsoventral expression differences across all genes within each subregion of each species. We observed significant positive correlations in all cases, supporting the biological relevance of the SVM-derived weights (Figure 2K).

We then investigated the similarity of dorsoventral gene expression patterns across different subregions. By comparing the correlation coefficients of SVM weights between subregions, we found that all subregions are positively correlated with each other (Figure 2L). This means that most genes with high weights toward the dorsal (or ventral) direction in one subregion also tend to exhibit high weights in the same direction in other subregions, suggesting a shared molecular architecture along the dorsoventral axis across the HPC and MEC. Notably, distinct intra-group patterns emerged, revealing subregion-specific similarities within the CA, DG, and MEC subregions, respectively (Figure 2L). In conclusion, our findings demonstrate conserved dorsoventral differences across species within all subregions of the HPC and MEC.

### Genetic properties characterizing the general dorsoventral differences across species

Next, we aimed to explore the conserved biological features underlying the dorsoventral gene expression patterns across species, as defined by the general SVM model. As previously demonstrated, high classification accuracy can be achieved and maintained across all subregions using the top 600 weighted genes (Figure 2I and J). To identify the key genes contributing to dorsoventral differences, we defined high-weight genes (HWGs) as those ranked among the top 300 most positively or negatively weighted genes for each subregion (Method). We also trained species-specific models and obtained the HWGs.

Figure 3 highlights the major functional modules enriched along the dorsoventral axis in CA, DG, and MEC with gene set enrichment analysis (GSEA)^43^. It also lists representative HWGs associated with these functions, as well as their dorsoventral weight rankings in both the general and species-specific SVM models. Detailed gene weights and expression differences are provided in Figure S6. Based on these results, we describe dorsoventral functional differences in CA, DG, and MEC from the perspective of nine enriched functional categories:

- Actin cytoskeleton dynamics. The dorsal CA region showed enrichment in actin cytoskeleton-related functions, including actin polymerization and depolymerization, actin filament network formation, and Arp2/3 complex-mediated actin nucleation. Several Arp2/3 complex subunit (ARPC) genes had high dorsal weights, particularly in CA1 and CA2 subregions. The actin cytoskeleton plays a pivotal role in neurons by regulating synapse formation, stabilization, and remodeling. Beyond shaping neuronal morphology, actin dynamics may also influence processes such as neuronal migration, dendritic branching, and axonal reconnection^44^. The dorsal CA region is likely to rely on this dynamic regulation to achieve more efficient neural network plasticity, thereby supporting complex cognitive functions, including learning, spatial information processing, and memory formation.
- MAPK signaling pathway. The dorsal CA and DG regions exhibited highly active signaling networks, particularly the classical mitogen-activated protein kinase (MAPK) pathway^45^. While both regions shared MAPK activity, the sets of HWGs differed. For instance, scaffolding genes such as KSR1^46^ and TRIB2^47^ had high dorsal weights in both regions, whereas dual specificity phosphatases (DUSP), such as DUSP5 and DUSP6^48^, were enriched only in the dorsal CA. PRKCB, a member of the protein kinase C (PRKC) family^49^, showed high weight in the dorsal DG. In summary, dorsal HPC cells have an adequate gene expression to rapidly respond to external stimuli through second messenger signaling, exhibiting a highly flexible regulatory network that provides a molecular basis for the execution of complex cognitive functions.
- Cell adhesion regulation. The dorsal CA region was enriched in various cell adhesion-related functions, including both cell-cell and cell-matrix adhesion, as well as their positive and negative regulation. Ephrin receptors from the EPH family^50^, such as EPHA7 and EPHA5, which guide cell migration and positioning, were highly weighted in the dorsal CA region. Through precise regulation of cell adhesion, the dorsal CA region can optimize the balance between neural network stability and plasticity.
- Neuronal development and myelination. Dorsal HPC, dorsal MEC, and ventral MEC all showed enrichment in neuronal development-related processes. Dorsal CA and DG were particularly enriched for axonogenesis, myelination, and dendritic development. During the morphological development of axons and dendrites, cytoskeletal components such as actin undergo continuous rearrangement and elongation, providing both structural support and driving force for axonal growth. This process is consistent with the observed enrichment of actin cytoskeleton dynamics in the dorsal CA region described above. In MEC, dorsal regions were enriched in axonal myelination, emphasizing the regulation of neural signal conduction speed and efficiency, with HWGs overlapping those identified in dorsal HPC. In particular, the myelin basic protein (MBP) gene exhibited high weights across all subregions of both the dorsal HPC and dorsal MEC. In contrast, ventral MEC was enriched in axon growth and extension, which are critical for establishing long-range neuronal connections and facilitating long-distance signal transmission, with genes such as FEZ1 and STMN2 showing high ventral weights. Semaphorin family genes^51^, such as SEMA7A, SEMA5B, and SEMA3C, were dorsal-enriched, while SEMA3A showed specific enrichment in ventral MEC, indicating regional specificity even for similar functions.
- Synaptic signaling and organization. Both HPC and MEC showed dorsoventral enrichment in synaptic functions, with region-specific emphasis. Dorsal HPC and ventral MEC were enriched in synaptic transmission, plasticity, and chemical signaling. In dorsal CA, glutamatergic signaling—particularly regulation of NMDA and AMPA receptors^52^—was prominent. For example, CNIH2, an AMPA receptor auxiliary protein, had high dorsal weight in CA. Ventral HPC and MEC were more focused on synaptic structure assembly. SYN2 had high ventral weights in all subregions of ventral HPC. Overall, the dorsal region predominantly emphasizes the regulation of excitatory signal transmission and the dynamic modulation of neurotransmitter activity, aligning with its role in fine-grained information processing, learning, and memory functions. In contrast, the ventral region places greater emphasis on the assembly and stabilization of synaptic structures, providing a stable foundation for neural network connectivity.
- Ion transport and homeostasis & calcium signaling. Dorsal HPC, especially the CA subregion, showed enrichment in ion—particularly calcium—transport and calcineurin signaling. Dorsal MEC showed positive regulation of calcium ion transport, with HWGs including CAMTA1, CALM1, and CAMKK2. Calcium transport and the calcineurin signaling play pivotal roles in modulating long-term potentiation (LTP) and long-term depression (LTD), thereby supporting the stabilization and updating of learning and memory. Ventral MEC was also enriched for calcium-related functions; however, it was more strongly associated with maintaining intracellular homeostasis, ion balance, and pH homeostasis, with HWGs like SLC4A10^53^ and ATP6V1A^54^.
- Cellular respiration and nucleotide metabolism. Dorsal MEC and ventral CA were enriched in aerobic respiration, oxidative phosphorylation, and electron transport chain activity, with more pronounced enrichment in ventral CA, potentially supporting higher metabolic activity and facilitating information storage. For example, CHCHD10 showed high dorsal weight only in dorsal MECint. Ventral CA showed enriched pathways in energy and purine nucleotide metabolism, especially synthesis, with genes such as AK5^55^ exhibiting high ventral weights. Mitochondrial complex I-related NADH:ubiquinone oxidoreductase subunits (NDUF)^56^ were differentially enriched: NDUFC2, NDUFA8, and NDUFB6 were ventral CA-enriched, while NDUFB9 and NDUFA1 were dorsal MEC-enriched.
- Behavior. Behavior-related gene signatures differed by region. Dorsal CA was enriched for cognitive functions such as learning and memory, while ventral DG was associated with fear response and social behaviors. In ventral DG, genes such as GRP^57^, CARTPT^58^, and CRHR1^59^ had high weights, indicating roles in emotion and social behavior regulation. These genes reflect the molecular distinctions along the dorsoventral axis of the HPC, providing a molecular basis for the region’s functional specialization.
- Amyloid regulation. Ventral MEC showed significant enrichment in amyloid-related processes, including metabolism of APP and the synthesis/clearance of amyloid-β (Aβ). HWGs in ventral MEC included APP and APOE. Aberrant APP processing leads to Aβ accumulation, a hallmark of Alzheimer’s disease (AD)^60^. The APOE ε4 allele is a major genetic risk factor for AD, promoting Aβ plaque formation^61^, suggesting a key role for ventral MEC in early AD susceptibility and pathology^62–64^.

**Figure 3:**
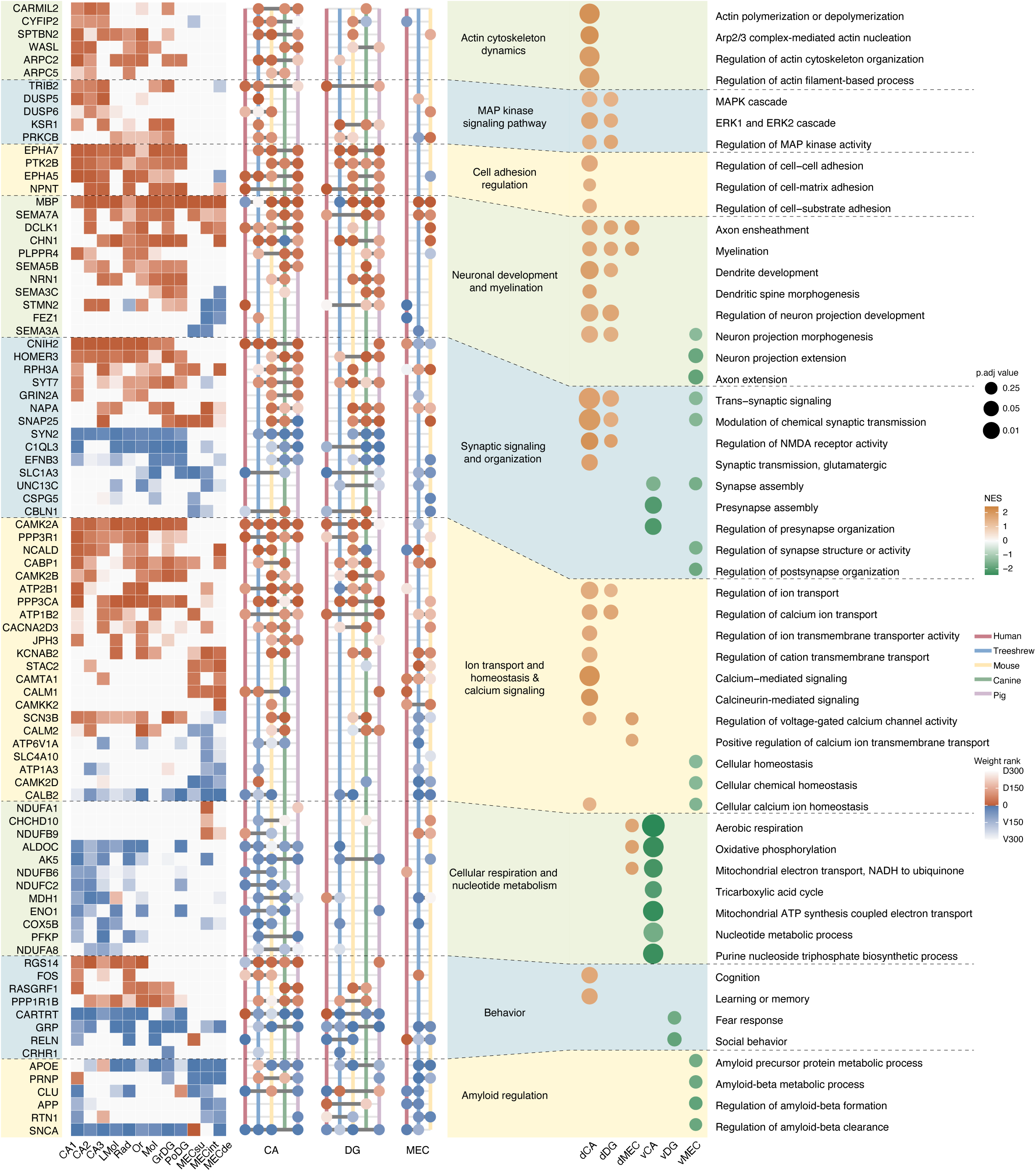
Dorsoventral functional modules and HWGs profiles across species and HPC/MEC subregions. Left: Heatmap showing the dorsoventral SVM weight ranks of selected HWGs in CA, DG, and MEC in the general model. The closer the rank is to 0, the higher the gene’s contribution to dorsoventral classification. Middle: dot plots showing the dorsoventral SVM weight ranks of selected HWGs in CA, DG, and MEC of five species-specific models. Ranks are normalized and averaged across subregions of CA, DG, and MEC, respectively. Ranks are then obtained according to the averaged weights. Each vertical line represents a species denoted by color. Dot colors represent the same as the left pannel. Right: Gene-to-function associations for each HWG, grouped into nine major functional modules. Dot plots showing gene ontology (GO) enrichment results from GSEA analysis of the dorsal and ventral compartments of CA, DG, and MEC (dCA/vCA, dDG/vDG, dMEC/vMEC). Dot color represents the normalized enrichment score (NES), and size corresponds to the Benjamini-Hochberg (BH)-adjusted p-values (two-sided permutation test) in GSEA.

### The major interspecies differences of HWGS reflecting divergent dorsoventral regulation across species

Considering the high accuracy of the general models in predicting combined data (Figure 2B), it seems intuitive that the dorsoventral differences identified in each species should be similar. Indeed, the gene weights for species-specific models positively correlated with the gene weights in the general model (Figure S7A), support this hypothesis.

However, we also noticed that the dorsoventral rankings of HWGs in the SVM general model showed variability when compared to their rankings in species-specific models. For instance, the gene CNIH2 exhibited high dorsal weights in the CA region across species-specific models constructed for the human, tree shrew, mouse, and pig. However, in the canine model, CNIH2 did not show a clear dorsoventral bias in weight distribution (Figure 3). Furthermore, even within species-specific models, the exact weight rankings of genes varied considerably. For example, ARPC2 ranked highly in the dorsal CA regions of tree shrew and mouse but ranked much lower in pig (Figure 3). In the MEC, interspecies variability in gene weights was even more pronounced. It was difficult to identify genes that consistently held high dorsal or ventral weights across all species (Figure 3). Additionally, some genes displayed reversed weight polarity between species. For example, RGS14 exhibited high dorsal weight in the CA region in the general model and in four out of five species-specific models, but in the human-specific model, it showed high ventral weight instead (Figure 3). To avoid potential biases introduced by pooling subregions within CA, DG, and MEC, we next examined the top-ranked HWGs identified by the general model and compared their weights to those from species-specific models for each subregion. Again, we observed substantial species-specific variability in HWG weights (Figure 4A).

**Figure 4:**
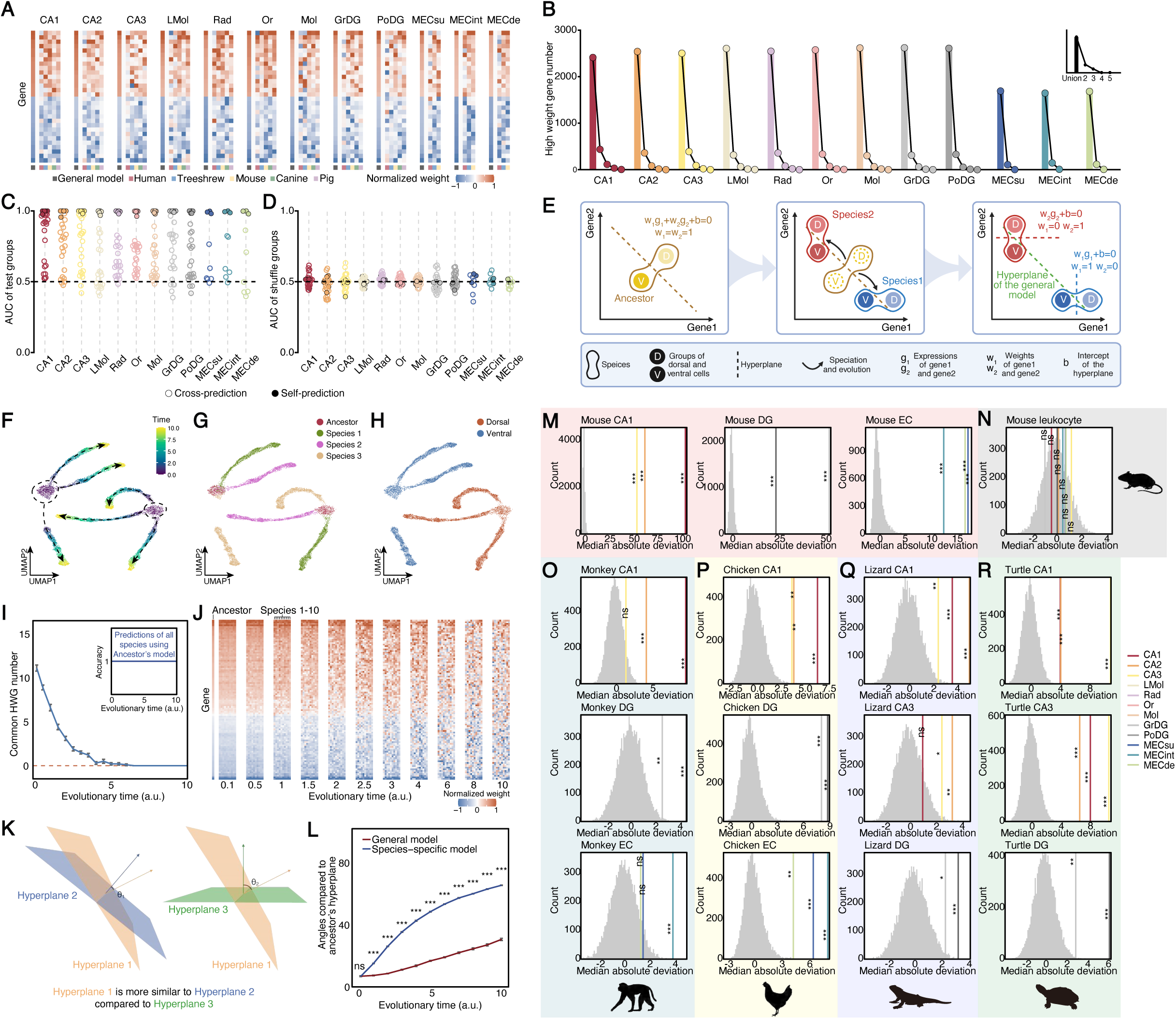
Species-specific dorsoventral differences and the ACT. (**A**) The weights of HWGs from the general model, as well as the corresponding weights in the species-specific models. Weights are normalized such that w_max_ = 1 and w_min_ = −1 for each subregion, while weights of 0 remain unchanged. (**B**) Common HWG number in n species. (**C**) AUC values for self-prediction (solid circles) and cross-predictions (hollow circles) of the test group in each subregion. (**D**) Same as in C for shuffle groups. (**E**) Schematic representation of the ancestral confinement theory (ACT). (**F**) UMAP representation of the simulation results of 3 species during evolution, labeled with evolutionary time (arbitrary unit). (**G**) Same as in F labeled with species. (**H**) Same as in F labeled with dorsoventral regions. (**I**) Simulation results showing the number of common HWGs across species during evolution. Error bars represent standard errors (SE) across 10 replicate simulations. The inset plot shows SVM accuracy at each evolutionary time point. (**J**) Simulation of the gene weights changes compared to ancestral gene weights during evolution. Weights are normalized such that wmax = 1 and wmin = −1 for each species, while weights of 0 remain unchanged. (**K**) Schematic representation of the hyperplane similarity defined by the angles. (**L**) Angles between the ancestral hyperplane and the hyperplanes of the general model or species-specific models. Error bars represent standard errors (SE) across 10 replicate simulations. Two-sided Wilcoxon tests were performed to compare angles. P-values are shown as star marks above the lines, where “* * *” represents p < 1e-3, “**” represents 1e-3 < p < 1e-2, “*” represents 1e-2 < p < 5e-2, and “ns” represents “non-significant”. (**M**) Projection of glutamatergic neuron expression data from the homologous HPC and MEC structures in chicken onto the normal vectors of the hyperplanes defined by the corresponding general models, compared to projections onto 10000 randomly generated directions. Variance is quantified using the median absolute deviation (MAD). One-sided permutation tests were conducted to assess significance, where “* * *” represents p < 1e-3, “**” represents 1e-3 < p < 1e-2, “*” represents 1e-2 < p < 5e-2, and “ns” represents “non-significant”. (**N**) Same as in (M) for lizard neurons. (**O**) Same as in (M) for turtle neurons. (**P**) Same as in (M) for rhesus macaque neurons. (**Q**) Same as in (M) for mouse leukocytes (negative control).

To further assess species specificity, we investigated the overlap of HWGs across species. Strikingly, the number of shared HWGs decreased rapidly as the number of species considered increased, with almost no HWGs shared across all five species in any subregion (Figure 4B). If HWGs are species-specific, it follows that dorsoventral differences in one species should also differ from those in another species. To verify this specificity, we cross-predicted the dorsoventral classification of one species using models trained on another species. Receiver operating characteristic (ROC) curves were generated, and the area under the curve (AUC) values were calculated. An AUC close to 1 indicates high prediction accuracy, whereas an AUC near 0.5 reflects no predictive power. The results revealed large variations in AUC across species, with some values close to 0.5 (i.e., no predictive power) (Figure 4C and D). In conclusion, these findings suggest that although the general SVM model captures molecular features that are conserved along the dorsoventral axis across species, substantial interspecies differences remain. Species-specific dorsoventral differences are evident, characterized by distinct sets of HWGs. This highlights the complexity and diversity of molecular regulatory mechanisms across species.

### Ancestral confinement theory explains the coexistence of the general and species-specific dorsoventral heterogenicities

To elucidate the phenomenon of dorsoventral differences being both conserved and species-specific, we propose the Ancestral Confinement Theory (ACT) (Figure 4E). According to ACT, a specific dorsoventral difference existed in the common ancestor of all species, representing a fundamental and essential feature that guided evolutionary trajectories and imposed selection pressures. Consequently, the ancestral hyperplane, which characterizes dorsoventral differences, remains relevant across all species, though it may not be optimal for each species due to evolutionary divergence (Figure 4E). This theory is supported by the evolutionary conservation of the HPC and MEC, or their homologous structures in for example the bird brain^65–67^, highlighting their conserved roles throughout evolution. A direct implication of ACT is that HWGs in one species may differ from HWGs in another species (Figure 4E). Moreover, ACT predicts that the dorsoventral heterogeneity defined by the general model (based on all species) is closer to the dorsoventral heterogeneity of the ancient ancestor compared to species-specific models. This general model reflects the most evolutionarily conserved features, representing the original dorsoventral differences.

Since obtaining gene expression profiles of the common ancestor is impossible, we perform numerical simulations to test the validity of ACT. First, we generate the gene expression profile of an ancestral species and define its dorsoventral hyperplane using SVM between dorsal and ventral cells. Next, we simulate evolutionary processes with confinement, applying genetic drift to the ancestor’s dorsal and ventral cells while ensuring they remain separable by the ancestral hyperplane. We use Brownian motion to simulate genetic drift^68,69^. While real-world selection mechanisms are more complex^70^, the specific mode of selection is not critical for ACT as long as confinement exists. For simplicity, we focus only on the impact of genetic drift, with further experimental data needed to validate ACT in the future.

UMAP shows that dorsoventral cells of all species diverge dramatically in their gene expression patterns (Figure 4F, G, and H) during the simulation. In contrast, all cells remain correctly classifiable by the ancestral hyperplane (Figure 4I, inset). However, the number of common HWGs across species rapidly declines to zero over evolutionary time (Figure 4I). This divergence is even more pronounced when examining the weights of all genes, as HWG profiles initially display similarity but diverge dramatically over time (Figure 4J), quite similar to the experimental observations (Figure 4A).

Next, we assess whether the general model is more similar to the ancestor’s model than species-specific models. Model similarity is evaluated by the angle between hyperplanes, with smaller angles indicating closer alignment (Figure 4K). We find that the angle between the general hyperplane and the ancestral hyperplane is significantly smaller than the angle for species-specific hyperplanes (Figure 4L). Furthermore, species-specific hyperplanes diverge from the ancestral hyperplane much faster than the general hyperplane as evolutionary time progresses (Figure 4L). In conclusion, ACT provides a framework to explain the observed experimental results. It predicts that the general model can effectively characterize the conserved dorsoventral heterogeneity of the common ancestor of the five species, despite species-specific divergences.

### Ancestral confinement suggests dorsoventral variations in HPC, EC, and amniote homologs

Next, we aimed to investigate whether dorsoventral differences also exist in the HPC and EC of non-human primates and even the evolutionarily conserved regions in non-mammalian species such as reptiles and birds. According to the ACT, dorsoventral differences defined by the general model approximate the ancestral characteristics conserved among the five mammalian species studied. Therefore, if a brain structure in another species exhibits gene expression patterns consistent with this dorsoventral heterogeneity, the gene expression values of its cells projected onto the normal vector of the general model’s hyperplane should display greater variance than projections onto random directions. This implies that the direction defined by the normal vector carries more biologically informative variation than random directions.

To test this, we obtained single-cell RNA sequencing data from the homologous HPC and EC structures of rhesus macaque (*Macaca mulatta*)^71^, chicken (*Gallus gallus*)^72^, lizard (*Pogona vitticeps*)^73^, and turtle (*Trachemys scripta elegans*)^73^. Only glutamatergic neurons were analyzed, as they constitute the majority of the neuronal population and exhibit clear subregion-specific distributions. For each species, cells from the CA region were evaluated using the general models of CA1, CA2, CA3, LMol, Rad, and Or; cells from the dentate gyrus (DG) were evaluated with general models of Mol, GrDG, and PoDG; and cells from the EC were evaluated with general models of MECsu, MECint, and MECde. As a negative control, we included single-cell RNA-seq data of mouse leukocytes^74^, which are not expected to show any dorsoventral differences. As a positive control, we used mouse HPC and EC single-cell data^75^. Strikingly, in all tested species, cells from each subregion showed significantly greater variance when projected onto the normal vectors of the corresponding general models compared to projections onto random directions (Figure 4M, O, P, Q, R). The positive control group showed consistent results (Figure S7B), whereas the negative control group exhibited no significant differences relative to random projections (Figure 4N). These findings suggest that dorsoventral heterogeneity is not restricted to mammals and may be a conserved evolutionary feature across amniotes.

### Phylogenetic analysis of dorsoventral expression profile reveals both conserved and divergent evolutionary trajectories

Since significant species-specific differences were observed, we further investigated how dorsoventral patterns differ among species and what genetic changes may underlie these evolutionary divergences. We first examined the phylogenetic relationships among the five studied species, using divergence times reported in a recent chronogram^76^. The most recent common ancestor of all species dates back to approximately 94 million years ago (MYA), with pig closely related to canine and human more closely related to tree shrew and mouse (Figure 5A). To assess species divergence from the perspective of dorsoventral gene expression, we constructed phylogenetic trees for each subregion based on SVM weight similarity, treating the general model as the ancestral root (Method). Surprisingly, the resulting trees often diverged from the known species phylogeny (Figure 5A). For instance, in CA1 and CA2, mouse, pig, and canine formed a cluster, whereas in CA3, human grouped more closely with pig and canine (Figure 5B and C).

**Figure 5:**
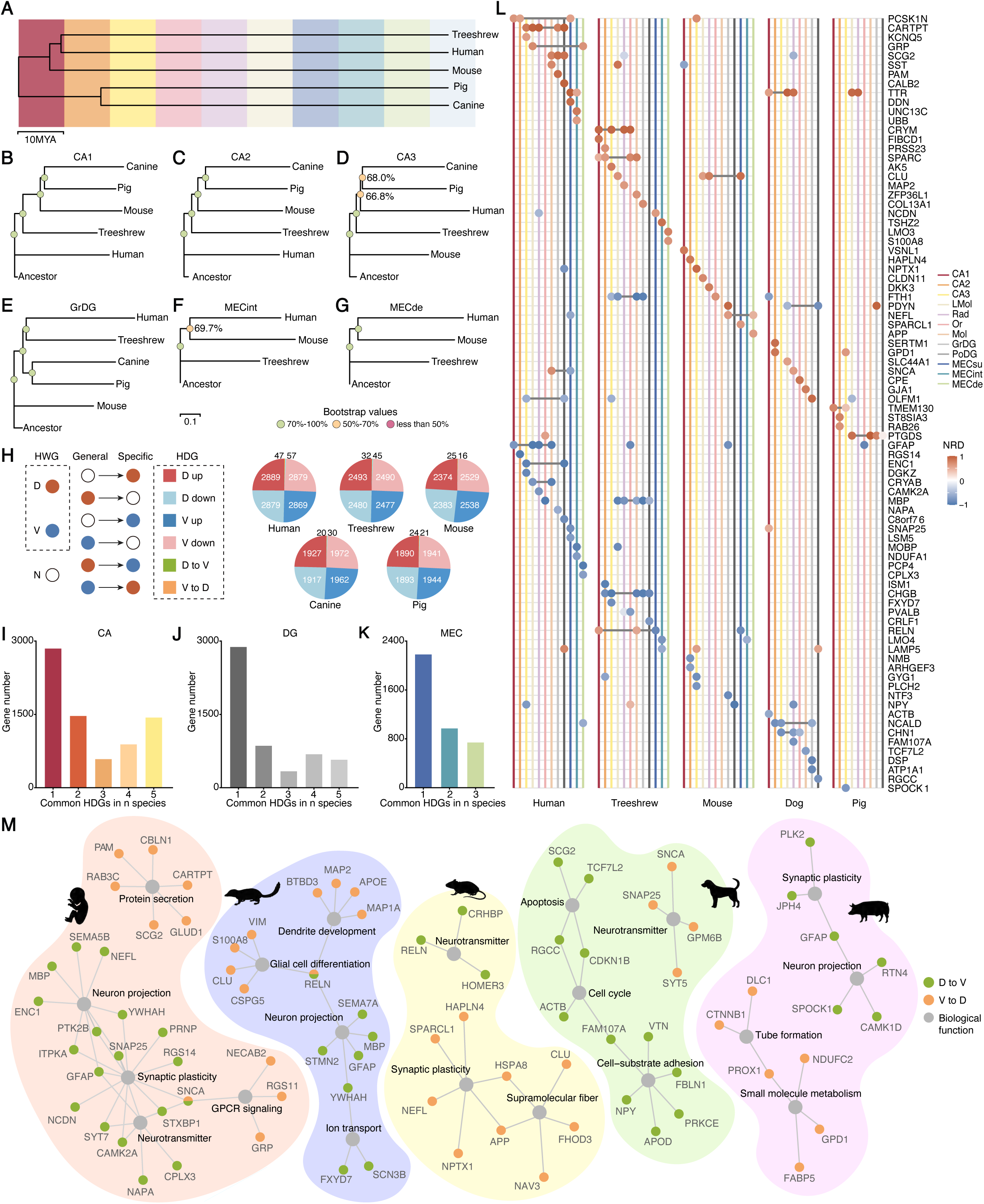
Species-specific changes during evolution. (**A**) Chronogram illustrating the evolutionary relationships among the five studied species. (**B**) Phylogenetic tree constructed based on the similarity of SVM weights between the general model (representing the ancestor) and species-specific models for CA1. Branch lengths are shown in arbitrary units. Bootstrap analysis was performed using resampling among orthologous genes to assess node support. Node colors indicate bootstrap support: green (> 70%), yellow (50̆70%), and red (< 50%). (**C**) Same as (B) for CA2. (**D**) Same as (B) for CA3. (**E**) Same as (B) for GrDG. (**F**) Same as (B) for MECint. (**G**) Same as (B) for MECde. (**H**) Schematic illustration of the six classes of HDGs (left) and their counts in each species (right). (**I**) Number of HDGs shared across n species in the CA region. (**J**) Same as (I) for DG. (**K**) Same as (I) for MEC. (**L**) Examples of reversed HDGs (“D to V” and “V to D”). Weight changes are quantified with normalized randk distance (NRD), where positive values indicate V-to-D shifts and negative values indicate D-to-V shifts. (M) GO functional enrichment analysis for the reversed HDGs.

Furthermore, the human CA3, GrDG, and MECde exhibited notably long branches, indicating accelerated evolution of dorsoventral features in these subregions (Figure 5D, E, and G). Interestingly, MEC subregions showed parallel patterns, with human clustering more closely with mouse than with tree shrew (Figure 5F and G). These findings suggest that subregional evolution in the HPC and MEC follows a species-independent trajectory.

To understand the molecular basis of these divergences, we identified High-Difference Genes (HDGs)—genes whose SVM weights in a given species and subregion significantly differ from those in the general model (Methods). HDGs were classified into six categories (Figure 5H): “D up” and “V up” (genes whose weights increase in the dorsal or ventral subregion), “D down” and “V down” (genes whose weights decrease in the dorsal or ventral subregion), and “D to V” and “V to D” (genes that reverse their weights completely, switching from dorsal to ventral HWGs or vice versa). Notably, the proportions of “D up”, “V up”, “D down”, and “V down” HDGs were relatively balanced across species, whereas “D to V” and “V to D” HDGs were substantially fewer. This finding suggests that most genes retain functional similarities in distinguishing the dorsoventral axis during evolution, with complete reversals being less favored, consistent with the conserved dorsoventral differences identified in the general model.

We then assessed how many HDGs were shared across species. The number of common HDGs across n species was calculated analogously to the common HWG metric (Method). Given the internal similarity among subregions within the CA, DG, and MEC domains, we performed this analysis separately for each domain. Interestingly, unlike HWGs, the number of common HDGs did not decrease sharply with increasing species count: HDGs both appeared exclusively in one species (i.e. species-specific HDGs) and were widely shared across multiple species (Figure 5I, J, and K). While species-specific HDGs reflected evolutionary divergence, the substantial set of HDGs shared across species suggests parallel divergence of functionally similar genes, potentially indicating convergent evolution.

To examine the most drastic molecular shifts, we focused on genes with complete weight reversals, i.e., “D to V” and “V to D” HDGs. To quantify these changes, we calculated the normalized rank distance (NRD), where positive values indicate a “V to D” reversal and negative values indicate “D to V”, with larger absolute values representing more substantial changes. Reversed HDGs displayed species-specific patterns across subregions (Figure 5L). In addition, when a gene was reversed in multiple subregions, the direction of reversal tended to be consistent. For instance, CARTPT reversed from ventral to dorsal in multiple human subregions, whereas CHGB showed the opposite pattern in tree shrew. CARTPT encodes a neuropeptide precursor involved in regulating appetite, energy balance, and stress responses, with links to addiction and obesity, while CHGB is a secretory protein in neuroendocrine cells that contributes to granule biogenesis and hormone release. Some genes, such as FTH1 and PDYN, reversed in opposite directions in different species, suggesting that their roles are highly context-dependent and shaped by species-specific pressures. FTH1 encodes the ferritin heavy chain, essential for iron storage and protection against oxidative stress. PDYN produces dynorphin peptides that act on κ-opioid receptors to modulate pain, stress, and addiction, with mutations linked to neuropsychiatric disorders.

To gain insight into the functional implications of reversed HDGs, we performed GO enrichment analyses for each species. Functions related to neural projections, plasticity, and neurotransmission were significantly enriched (Figure 5M). Notably, these functions sometimes evolved in opposite directions: plasticity-related genes were reversed from dorsal to ventral in human and pig but from ventral to dorsal in mouse. These findings suggest that while the general functional categories are conserved, the molecular mechanisms supporting them diverge across species to meet specific ecological or behavioral demands. Interestingly, projection- and plasticity-related functions were also conserved in the general model (Figure 3), suggesting that functional outcomes are preserved through different gene sets in different species—a hallmark of evolutionary convergence. Thus, the evolution of weight-reversed genes reveals both divergence and convergence in the genetic architecture underlying dorsoventral patterning.

### Consensus cell types across five species demonstrate conserved dorsoventral cell-type distributions, with region-specific excitatory and broadly dispersed inhibitory neurons

Next, we investigated whether common cell type differences exist along the dorsoventral axis across species. To address this, we established a consensus single-cell reference for the hippocampal formation (HPF, including HPC and EC) in mouse, tree shrew, pig, canine, and human. First, we generated single-nucleus RNA sequencing data for the tree shrew HPC and MEC. Additionally, we collected single-cell RNA sequencing data for the mouse HPC and MEC^75^ [75] and single-nucleus RNA sequencing data for the pig HPC^77,78^, canine HPC^79^, and human HPC and MEC^80^ from public repositories. After normalization and integration (Methods), we employed graph-based clustering to identify 58 clusters, which are further grouped into 37 metaclasses using hierarchical clustering (Figure 6A, B, and C). UMAP representations show that cells from different species are effectively integrated (Figure 6A and B). Common markers for these clusters are identified (Figure 6C). In the dendrogram of all cell types, neurons and non-neurons are hierarchically separated. Neurons are further divided into excitatory and inhibitory subtypes, except for one type, “Glut HPF CR” (Figure 6C). Excitatory neurons were named according to their primary anatomical regions, except for “Glut Mix” 1-5, which include multiple classes of neurons, such as “splatter” neurons in human and some excitatory neurons in other species. Inhibitory neurons were classified based on their markers, except for “GABA Mix CGE” 1 and 2, which include subsets of NDNF and VIP GABAergic neurons. These cell types correspond well to those previously defined in mouse and human studies^75,80 80^ (Figure S8).

**Figure 6:**
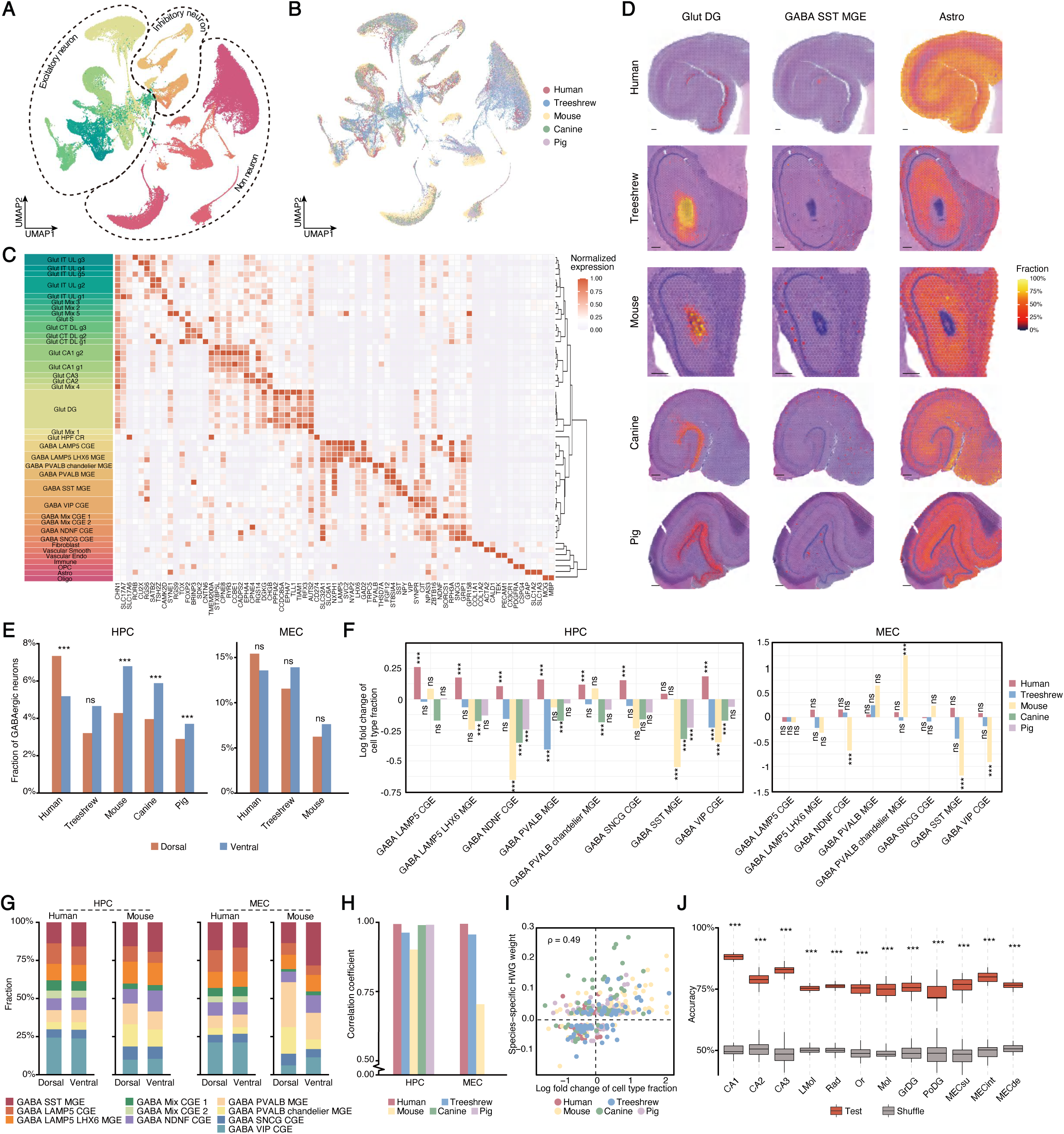
Consensual cell types and cell type differences across species. Abbreviations: Glut, glutamatergic; HPF, hippocampal formation; CR, Cajal-Retzius; GABA, GABAergic; CGE, caudal ganglionic eminence; MGE, medial ganglionic eminence; Vascular Smooth, smooth muscle cell; Vascular Endo, endothelial cell; Astro, astrocyte; Oligo, oligodendrocyte; OPC, oligodendrocyte precursor cell. (**A**) UMAP representation of identified cell types. (**B**) UMAP representation with species labeled. (**C**) Left panel: Markers for all 37 metaclasses of cell types. Cell type colors are the same as in (A). Right panel: hierarchical clustering of all cell clusters. (**D**) Spatial distributions of selected cell types across different species. (**E**) Proportional differences in overall GABAergic neurons along the dorsoventral axis across species in HPC (left) and MEC (right). Cellular composition data were first normalized using centered log-ratio (CLR) transformation, which were then pooled by averaging the fraction in HPC or MEC. Two-sided Wilcoxon tests were used for statistical comparison, and p-values were adjusted using the BH method. To reduce the false-positive rate, an additional threshold based on the magnitude of proportional differences was applied to determine statistical significance (Method), where “* * *” represents statistically significant differences, and “ns” represents “non-significant”. (**F**) Dorsoventral differences in distinct metaclass of GABAergic neurons in HPC (left) and MEC (right) across species. Log fold change represents the difference between the normalized proportions of each GABAergic neuron metaclass in dorsal versus ventral regions. Positive values indicate higher proportions in the dorsal region, while negative values indicate higher proportions in the ventral region. (**G**) Proportions of GABAergic neuron metaclasses within total GABAergic populations along the dorsoventral axis of HPC (left) and MEC (right) in human and mouse. (**H**) Pearson correlation coefficients of the metaclass proportions between dorsal and ventral regions across species in both HPC and MEC. (**I**) Correlation between the log-transformed differences in cell type proportions and the SVM weight values of their corresponding marker HWGs in species-specific models. The Pearson correlation coefficient (ρ) is shown in the upper left corner of each plot. Two-sided t-tests are performed (p = 4.41e-21). (**J**) SVM classification accuracies for each subregion. Models were trained and tested using CL2-transformed cell type compositions from all five species (for MEC, data from only human, mouse, and tree shrew were used). Two-sided Wilcoxon tests compared the accuracy between the testing and shuffled groups. P-values were corrected using FDR, with significance levels indicated by asterisks, where “* * *” represents FDR < 1e-3.

To validate the defined cell types and analyze their spatial distribution, we used Cell2location^81^ for deconvolution of the spatial transcriptomics data with the defined cell types (Methods). We observed that the spatial distribution of cell types is highly conserved across species. Excitatory neurons localized well to their expected spatial regions, while inhibitory neurons showed a sparse distribution (Figure 6D). Astrocytes were enriched in subregions dense in axons and dendrites, consistent with their role in neuronal support (Figure 6D). Together, these results establish a cross-species cell type reference of the HPC-MEC network, providing a framework to explore dorsoventral molecular and cellular programs in evolution.

Unlike the widespread dorsal enrichment of GABAergic neuron metaclasses in the human HPC, the other species showed more metaclass-specific dorsoventral distribution patterns (Figure 6E and F). Further analysis of each metaclass’s proportion within the total GABAergic population revealed considerable dorsoventral differences in mouse, particularly in HPC (Figure 6G), with the lowest dorsal-ventral correlation among all species (Figure 6H), indicating a more metaclass-dependent distribution in mouse.

Although overall GABAergic neuron proportions in the MEC did not differ significantly between dorsal and ventral regions (Figure 6E), specific metaclasses still exhibited significant differences.

In mouse, the same metaclasses enriched ventrally in the HPC—GABA SST MGE, GABA VIP CGE, and GABA NDNF CGE—were also enriched in the ventral MEC. In contrast, the GABA PVALB chandelier MGE metaclass was significantly enriched in the dorsal MEC (Figure 6F). This metaclass targets the axon initial segment of pyramidal neurons and is critical for precise spatiotemporal control of action potential firing. In comparison, no significant dorsoventral differences in GABAergic metaclass distributions were observed in the MEC of human and tree shrew (Figure 6F). Analysis of GABAergic metaclass composition within the total GABA population in MEC showed that mouse exhibited the most pronounced dorsoventral variation (Figure 6G), with dorsal-ventral correlations much lower than those in human and tree shrew (Figure 6H). Furthermore, in mouse, the magnitude of dorsoventral differences in both overall cell proportion and relative metaclass composition of GABAergic neurons was greater in MEC than in HPC, indicating stronger dorsoventral heterogeneity in the MEC.

In summary, the result demonstrates pronounced species-specific patterns in the dorsoventral distribution of GABAergic neurons, with varying trends across brain regions. In the human HPC, a broad increase in GABAergic neurons in the dorsal region was observed, with all metaclasses contributing modestly. In contrast, the ventral enrichment of GABAergic neurons in the HPC of other species was largely driven by specific metaclasses—particularly GABA SST MGE, GABA VIP CGE, and GABA NDNF CGE. In the MEC, dorsoventral differences were most prominent in mouse, where metaclass distribution trends mirrored those in the HPC, suggesting potential functional coordination between the two regions in emotional regulation, stress responses, and behavior modulation.

### Integration of cell-type composition and SVM modeling reveals conserved and species-specific dorsoventral patterning

To further validate the reliability of dorsoventral distribution across cell types, we integrated cell-type composition data with gene weight information derived from SVM models. Theoretically, if a particular cell type is enriched in the dorsal (or ventral) region, its corresponding marker genes should exhibit higher expression levels in the same region. Accordingly, those genes should also exhibit higher weights in the SVM model. We extracted weights from each species-specific SVM model and calculated the correlation between the dorsoventral distribution of each cell type and the weights of their corresponding HWGs. The analysis revealed a significant positive correlation (Figure 6I), indicating that marker genes assigned higher weights in the SVM model tend to be associated with cell types that exhibit pronounced bias in dorsoventral distribution.

Building upon this observation of interspecies differences in cell-type distributions and considering the high accuracy of the SVM model in identifying dorsoventral differences—as well as the strong alignment between marker gene weights and cell-type spatial distributions—we further tested whether dorsoventral classification could be achieved directly using cell-type composition alone. To this end, we compiled dorsoventral cell-type proportion data across all species and trained a general SVM model based solely on these cell composition features. The results showed that the classification accuracy of this cell-type-based SVM model was significantly higher than that of the shuffled control (Figure 6J), suggesting that dorsoventral information is indeed embedded within the cellular composition and that this pattern exhibits a degree of cross-species conservation.

## Discussion

Our cross-species spatial transcriptomic analysis reveals a conserved yet evolutionarily flexible molecular architecture along the dorsoventral axis of the HPC and MEC, offering unprecedented insights into how these regions balance functional specialization with evolutionary adaptation. By profiling five phylogenetically diverse species—tree shrew, mouse, pig, canine, and human—we demonstrate that dorsoventral heterogeneity is governed by a core set of evolutionarily constrained gene network, while accommodating species-specific adaptations through divergent molecular mechanisms.

### Construction of a Multispecies Dorsoventral Molecular Atlas of the HPC and MEC

In this study, we applied spatial transcriptomics to HPC and MEC of five mammalian species—human, tree shrew, mouse, canine, and pig—and, for the first time, generated a cross-species molecular atlas along the dorsoventral axis. Leveraging the high spatial resolution of 10X Visium, we precisely delineated internal subregions of the HPC (cell body enriched regions CA1, CA2, CA3, GrDG, PoDG; and neuropil enriched regions Or, Lmol, Rad, Mol) and of the MEC (MECsu, MECint, MECde), annotated each as dorsal or ventral, and systematically characterized their molecular signatures.

Although spatial transcriptomics achieves near single-cell resolution, it does not attain true single-cell granularity, limiting precise cell-type annotation. To mitigate this, we integrated spatial transcriptomic data with single-cell (or single-nucleus) RNA-seq profiles and applied a reference-based deconvolution algorithm to infer cellular composition. While this strategy markedly improves cellular annotation, some prediction bias persists. Future incorporation of ultra-high-resolution imaging modalities (e.g., Stereo-seq, MERFISH)^82,83^ will be essential to further refine cell-type assignment accuracy.

### Conserved Molecular and Cellular Signatures of Dorsoventral Organization

Despite ∼94 million years of evolutionary divergence, all five species exhibited striking conservation of dorsoventral gene expression gradients in HPC and MEC subregions. Supervised machine learning (SVM and LDA) classified dorsal and ventral spots with high accuracy (Figure 2B), even in white-matter-rich areas, suggesting that dorsoventral patterning is a fundamental organizing principle. This conservation extended to functional gene modules: the Dorsal HPC prioritized actin cytoskeleton dynamics (e.g., ARPC2/3), MAPK signaling (e.g., KSR1, DUSP5), and synaptic plasticity (e.g., CNIH2), aligning with their roles in high-precision spatial coding and memory^84^. The Ventral HPC/MEC enriched amyloid regulation (e.g., APP, APOE), social behavior genes (e.g., CARTPT), and synaptic stabilization (e.g., SYN2), consistent with its involvement in emotional processing and early Alzheimer’s disease (AD) pathology^62–64^. Notably, human and tree shrew exhibited the greatest number of reverse genes (Figure 5H), implying accelerated evolution of their dorsoventral molecular landscapes. Particularly, multiple neuropeptide-related genes shifted from a ventral to a dorsal bias in the human HPC, suggesting an expansion of emotion- and social-behavior-related functions in the dorsal region. Given the richer emotional repertoire in human, this functional extension may stem from remodeling of axis-specific expression patterns at the molecular level.

The cellular deconvolution linked these molecular gradients to conserved dorsoventral distributions of inhibitory neuron subtypes (Figure 6). For instance, GABAergic PV+ cells were dorsally enriched in mouse MEC—a potential substrate for theta rhythm generation^85,86^—while SST+ and VIP+ interneurons dominated ventrally, mirroring their roles in affective processing^87–89^. This conservation suggests that dorsoventral specialization arises from a shared developmental blueprint, reinforced by selective pressures. In contrast, in the human HPC—apart from SST-MGE cells—nearly all GABAergic subtypes exhibited modest yet significant dorsal enrichment. This shift may reflect an evolutionary elaboration of the dorsal human HPC’s involvement in higher-order cognition and memory, as well as its emerging role in emotional control, necessitating more precise and potent inhibitory regulation to maintain network stability and information processing efficiency.

### Evolutionary Divergence and the Ancestral Confinement Theory (ACT)

While dorsoventral organization was broadly conserved, high-weight genes (HWGs) defining these axes showed remarkable species specificity (Figure 4). For example, CNIH2, a dorsal AMPA receptor regulator in most species, lost dorsoventral bias in canine, RGS14 switched from dorsal (mouse) to ventral (human) weighting, implicating divergent roles in cognition versus emotion. Notably, multiple neuropeptide-related genes shifted from a ventral to a dorsal bias in the human HPC, suggesting an expansion of emotion- and social-behavior-related functions in the dorsal region^90^. Given the richer emotional repertoire in human, this functional extension may stem from remodeling of axis-specific expression patterns at the molecular level.

We proposed the ancestral confinement theory (ACT, Figure 4), which allows genetic divergence while preserving functional topography. Our ACT theory suggests the conserved dorsoventral axis from reptiles to mammals indicates an origin in a common amniote ancestor. It indicates the functional segregation between spatial and affective processing provided key adaptive value. Cross-species and simulation studies confirm that this systems-level organization persists despite genetic divergence, illustrating a classic case of systems-level convergent evolution^91^, where diverse molecular mechanisms achieve similar functional outcomes under shared constraints. ACT identifies developmental patterning as the core evolutionary constraint, consistent with developmental bias^92^. Conserved morphogen gradients, transcription factors, and connectivity rules form a hyperplane that ensures dorsoventral polarity, enabling a many-to-one genotype-to-phenotype mapping^93^.

Our findings illuminate how dorsoventral specialization supports region-specific computation while conferring differential disease vulnerabilities. This dorsoventral specialization can be related to a cognitive affective tradeoff. By integrating spatial transcriptomics across five species, we unveil a unifying yet flexible blueprint for dorsoventral organization in the HPC-MEC network. Evolutionary constraints preserve core functional axes, while genetic drift enables species-specific adaptations—a duality elegantly explained by ACT. These insights not only advance our understanding of hippocampal evolution but also provide a roadmap for studying region-specific vulnerabilities in neuropsychiatric disorders, such as aging, emotion and neural degeneration related cognitive decline.

## Methods

### Experimental materials

The collection and study protocol for de-identified human fetal brain tissue was approved by the Biomedical Ethics Committee of Peking University (approval No. IRB00001052-24193). All samples were obtained from donors who had voluntarily elected to terminate pregnancy and had provided written informed consent in full compliance with national laws and institutional ethical regulations governing the use of such tissues. The protocol received formal approval from the Biomedical Research Ethics Committee for use of postmortem fetal tissue. All procedures were conducted in accordance with the Administrative Regulations on Human Genetic Resources issued by the Ministry of Science and Technology of China.

All procedures involving animals were approved by the Institutional Animal Care and Use Committee (approval No. LSC-MiaoCL-4) and were conducted in accordance with relevant ethical guidelines. Experimental mice were of the C57BL/6 strain and were housed in a temperature-controlled (22-24 ℃), ventilated facility, with no more than five mice per cage. A 12-hour light/dark cycle was maintained (lights off at 8:00, lights on at 20:00), and all experiments were conducted during the dark phase. Tree shrew was singly housed in similarly controlled and ventilated environments, with the same 12-hour light/dark cycle. Fresh brain tissues from canine (beagle canine) and domestic pig (Duroc Landrace Large White hybrid) were obtained directly from collaborating institutions and did not require on-site housing.

### Preparation of spatial and single-nucleus transcriptomic samples

Spatial transcriptomics was performed using the 10x Genomics Visium platform, including both the standard Visium Spatial Gene Expression (PN-1000184) and the high-resolution Visium HD (PN-1000675) version. Fresh-frozen brain tissues from tree shrew, mouse, canine, and pig were processed for the standard Visium assay, while formalin-fixed paraffin-embedded (FFPE) human fetal brain samples were analyzed using Visium HD.

For fresh-frozen samples, brain tissues were embedded in OCT and sectioned sagittally at 10 µm thickness using a cryostat (−10°C). Sections were mounted onto Visium slides, ensuring coverage of the hippocampal (HPC) and medial entorhinal cortex (MEC) dorsoventral axis. For human fetal brain, tissues were fixed in 4% paraformaldehyde at 4°C for 72 hours, sectioned into ∼5 mm slabs, and further processed for paraffin embedding. FFPE blocks were cut into 5 µm sections, floated on a water bath, and mounted on Visium HD slides. For both tissue types, anterior (ventral) and posterior (dorsal) regions of the HPC axis were sampled.

On the Visium slide, mRNA capture relies on spatially barcoded oligonucleotides. The standard Visium platform achieves an average resolution of 1–10 cells per spot (55 µm diameter, ∼5,000 spots per capture area), whereas Visium HD enables subcellular resolution with ∼11 million 2 × 2 µm² barcoded squares per capture area.

Tissue optimization (TO) experiments were conducted following 10x Genomics protocols (CG000238) to determine optimal permeabilization times: 22 min for pig and 18 min for tree shrew, mouse, and canine brain tissues. Subsequent sample processing followed the manufacturer’s user guide (CG000239 for Visium; CG000685 for Visium HD). Briefly, sections were methanol-fixed, H&E-stained for histological alignment, and imaged prior to mRNA release and capture. In the Visium HD workflow, FFPE sections underwent probe hybridization and processing with the CytAssist instrument for precise alignment of tissue and capture areas. Captured transcripts were reverse-transcribed to cDNA, followed by library construction and quality control. Sequencing was performed on Illumina NovaSeq using PE150 mode, with 125 million reads per capture area for Visium and 275 million reads per capture area for Visium HD.

Single-nucleus RNA-seq was performed on tree shrew brain tissues using the 10x Genomics Chromium system (PN-1000494) and following the Chromium Nuclei Isolation protocol (CG000505). Fresh-frozen 50 µm sections covering HPC and MEC were manually dissected, homogenized in lysis buffer, and filtered. Nuclei were purified, DAPI-stained, and sorted by flow cytometry. Approximately 35,000 nuclei per sample were used for library preparation. Sequencing was carried out on Illumina NovaSeq at 25,000 PE150 reads per nucleus.

## Software

Most analyses were conducted in R (v4.0.1). Python (v3.9) was used for deconvolution of cell types, and R (v4.2.2) was employed for functional enrichment analysis.

## Code availability

All codes for data analysis in this study are available at https://github.com/QirunWang/Codes_for_dorsoventral_analysis.

## Data availability

All raw and processed data generated in this study will be made publicly available upon publication.

Human snRNA-seq data in the format of read count matrices^80^ were retrieved from Cellxgene (https://cellxgene.cziscience.com/collections/283d65eb-dd53-496d-adb7-7570c7caa443), including tissues from the HPC and EC. These data were used to construct the consensus cell type atlas across species.

Human snRNA-seq data in the format of read count matrices^26^ were retrieved from the NCBI GEO database (GSE160189). Metadata for the experiments were obtained from (https://cells.ucsc.edu/human-hippo-axis/meta.tsv). These data were used to validate our general models.

Mouse scRNA-seq data in the format of read count matrices and metadata^75^ were retrieved as described in their published paper. Only the “10xv3” dataset was used. Cells from “HPF” (hippocampal formation) samples were selected, and supertypes not present in the HPC or EC in the MERFISH dataset^94^ were filtered out.

Pig snRNA-seq data for the HPC, including raw sequencing reads and metadata^78^, were retrieved from the NCBI GEO database (GSE186538).

Canine snRNA-seq data in the format of read count matrices and metadata^79^ were requested directly from the authors. Chicken snRNA-seq data for brain in the format of read count matrices and metadata^72^, were retrieved from heiData^72^.

Lizard and Turtle scRNA-seq data for brain in the format of read count matrices and metadata^73^, were retrieved as described in their published paper.

Rhesus macaque snRNA-seq data for HPC and MEC in the format of read count matrices and metadata^71^ were retrieved from Cellxgene (https://cellxgene.cziscience.com/collections/8c4bcf0d-b4df-45c7-888c-74fb0013e9e7).

Mouse scRNA-seq data for immune cells in the format of read count matrices and metadata^74^ were retrieved from Cellxgene (https://cellxgene.cziscience.com/collections/7e216a15-82df-46ee-b454-d0261d99e5f5).

Phylogenetic relationships of the five species were retrieved from TimeTree^76^.

## Data alignment and cleaning

Raw sequencing reads were processed using Spaceranger (v2.1.1) for spatial transcriptomes and Cellranger (v7.2.0) for single-nucleus transcriptomes, with default parameters. Reads from mouse and human were mapped to 10X prebuilt genome references mm10 - 2020 - A and GRCh38 - 2020 - A, respectively. For other species, genome references were downloaded and built following the tutorials provided by Spaceranger and Cellranger. For pig, genome reference Sscrofa11.1 (GCF_000003025.6) was downloaded from NCBI RefSeq. For canine, genome reference ROS_Cfam_1.0 (GCA_014441545.1) was downloaded from ENSEMBL release 113. For tree shrew, genome reference TupChi_ 1. 0 (GCF_000334495.1) was downloaded from NCBI RefSeq.

The output read matrices (and image data for spatial transcriptomes) were analyzed using the Seurat package (v4.0.2)^40^ in R. Gene names were standardized to ENSEMBL IDs to avoid duplicated symbols. Mitochondrial genes were excluded from the analysis. For snRNA-seq data, additional quality control filters were applied: cells with fewer than 500 counts, more than 3% mitochondrial counts, or more than 25,000 counts were removed to exclude low-quality cells and doublets.

## Anatomic subregion registration

For human, mouse, and tree shrew, HPC and MEC subregions were delineated based on anatomical landmarks defined in their respective brain atlases. For pig and canine, where these regions are less well characterized, subregion registration was guided by both anatomical features and the spatial patterns of unsupervised spot clustering. Notably, we did not distinguish the CA4 (Cornu Ammonis 4) region, as studies suggest CA4 is not distinct from PoDG in terms of cellular composition^38,39^.

## Spot binning for human data

The smallest bin sizes are 2 μm squares for 10 X Visium HD and 55 μm circles for 10 X Visium. To homogenize the data for subsequent combined analyses, we binned the human HD data into 48 μm squares, ensuring their area (2304 μm^2^) is comparable to that of the non-HD spots (2375 μm^2^). The 48 μm-binned spots were used for all subsequent analyses.

## Data normalization, scaling, and batch correction

We applied SCTransform (SCT)^95^ for normalization, scaling, and batch correction of the data. Samples from the same experimental batch were merged before running SCT. SCT calculates Pearson residuals based on the background distribution of gene expression across all spots within a batch, making it critical to maintain a consistent background distribution for each sample. To ensure consistency of the distribution, we selected spots belonging to the following anatomical structures: HPC (including CA, DG, subiculum, alveus (alv), and fimbria (fi)), MEC, and lateral entorhinal cortex (LEC), ensuring that every SCT batch contained all these regions. For pig and canine samples, where MEC data were unavailable, we included spots from the cortex to compensate for the missing regions. To accelerate the SCT process, we used glmGamPoi (v1.2.0). The scaled data output from SCT was used for all subsequent analyses.

## Obtaining orthologous genes across species

Orthologous genes between human and other species were retrieved from NCBI using the NCBI Datasets command-line tools (v2.20.0)^96^ . Only one-to-one mappings of orthologous genes were retained (Figure S4A and B). For consistency in subsequent analyses, the gene names of all non-human species were renamed to their corresponding human gene names. For chicken data from^72^, due to an old version of the genome reference (galGal5) and ENSEMBL ID was used, we got the orthologous genes of human directly from the supplementary table of the reference^97^.

## Integration of data from different species

To minimize species-specific variations, we applied Seurat Integration^98^ across datasets (Figure 1B and C, Figure 6A and B).

For spatial transcriptomic data, 3,000 anchor features were selected, and canonical correlation analysis (CCA) reductions were used in the integration pipeline.

For sc/snRNA-seq data, we first downsampled the total number of cells in each species. Human and mouse data were randomly downsampled to 20k cells, preserving the original proportions of annotated cell types. Tree shrew data were randomly downsampled to 30k cells. Pig and canine data were randomly downsampled to 10k cells, as only HPC data were available for these species. The downsampled cells were then integrated using the same parameters as the spatial transcriptomes, with one exception: reference datasets were specified during integration. For each species, the dataset with the largest number of cells was designated as the reference.

The resulting integrated data were used exclusively for Principal Component Analysis (PCA), Uniform Manifold Approximation and Projection (UMAP) visualization, clustering, and cell type annotation, which are also done in Seurat.

## Hierarchical clustering

Hierarchical clustering was performed to compare different groups of spots (Figure 1D) or cells (Figure 6C). Using the integrated data, average PCA values were first calculated for each group. Distances between groups were then computed as 1 - corr(averagePCA), where corr represents the Pearson correlation coefficient between groups. Finally, hierarchical clustering was performed using the complete linkage method based on the calculated distances.

## Finding gene markers

Markers for subregions (Figure 1D, Figure S4F) or cell types (Figure 6C) were identified using FindMarkers by comparing the target spots/cells to the remaining spots/cells. The slot parameter was set to “scale.data”, and test.use was set to “wilcox”. Markers were further filtered using a 0.05 FDR cutoff.

## Supporting vector machine (SVM) analysis

We utilized SVM to identify general and species-specific dorsoventral differences (Figure 2). Models were trained using either data from all species (general models) or data from each species individually (species-specific models). For the general model in each subregion, spots were randomly sampled to ensure equal numbers of dorsal and ventral spots and an equal number of spots from each species. For the species-specific model in each subregion, only the number of dorsal and ventral spots was balanced. Data with shuffled dorsoventral tags were generated from the randomly sampled data to serve as controls. SVM computations were performed using the R package e1071 (v1 .7- 9) with the following parameters: type set to “C-classification”, scale set to “False”, and kernel set to “linear”. For each model, 80% of the data were used for training and 20% for testing. Each process was replicated 20 times. After training, the weights of genes and intercepts were extracted, L2 normalized, and averaged across all replicates.

We also trained general models with the top n weighted genes removed (Figure 2I), or using only the top n weighted genes (Figure 2J) for each subregion. SVM weights contain both positive and negative values, with genes having weights far from 0 (i.e., large absolute weights) being more important for classification. Genes were ranked in descending order based on their absolute average weights for each subregion. Models were then trained using expression matrices containing only the top-n weighted genes, or without top n weighted genes, and their performance was tested. Each model was replicated 5 times. To visualize the resulting accuracies, the data were smoothed using ksmooth with a normal kernel and bandwidth ranging from 50 to 1000 before being plotted.

Dimensionality reduction was performed to visualize the classification results of the trained SVM models (Figure 2B). Axis 1 corresponds to the direction of the normal vector of the SVM hyperplane, while axis 2 represents an arbitrary orthogonal direction. Data points were projected onto the two-dimensional plane defined by these axes. The projection was then centered along Axis 1 so that the SVM decision boundary aligns with x = 0.

For general SVM models trained on cell type data (Figure 6J), CLR-transformed compositions were used as input. All processes followed the same steps as described above.

## Linear discriminant analysis (LDA)

We utilized LDA to validate the general dorsoventral differences identified by SVM (Figure S5F and G). The R package MASS (v7.3-51.6) was used for training with default parameters. All procedures for LDA followed the same steps as those used for the general SVM model.

## SVM distance analysis

SVM distance is defined as the shortest distance between a spot and its SVM hyperplane, determined by the average weights of the genes and the average intercepts (Figure 2C, D, E, F, G, H, and Figure S5H). For anatomic distance, in tree shrew and mouse, this is defined as the vertical coordinate of a spot on the sagittal section. The anatomic distances of all spots on the same section are then normalized to a range of -1 to 1. Finally, Pearson correlation coefficients are calculated between the normalized anatomic distance and the SVM distance.

## Correlation between the SVM weights and expression

Pearson correlation coefficients were calculated between the expression levels and the SVM weights across all genes for a given subregion (Figure 2K). The expression level of a gene was defined as the average difference of the SCTransformed scaled expression along the dorsoventral axis within the subregion. The weight of a gene was taken as the average weight from the general SVM model for that subregion.

## Gene set enrichment analysis (GSEA)

We performed GSEA on genes in CA, DG, and MEC (Figure 3). SVM weights were first L2-normalied and averaged across subregions in CA, DG, and MEC. Genes were then ordered by the SVM weights. GSEA was then performed using the GO database^99,100^ in R (v4.2.2). The gseGO function of R package clusterProfiler (v4.6.2)^101^ was used. The cut-off criteria were set as the p-value < 0.05 and the false discovery rate (FDR) q value < 0.25 . The enriched terms were manually reviewed and displayed in the plots.

## Gene ontology (GO) functional enrichment analysis

We performed functional enrichment analysis for the reversed HDGs (Figure 5M) in each species using the GO database^99,100^ in R (v4.2.2). The enrichGO function from the R package clusterProfiler (v4.6.2)^101^ was used. The cut-off criteria were set as the p-value < 0. 05 and the false discovery rate (FDR) q value < 0.25 . The enriched terms were manually reviewed and displayed in the plots.

## High-weight gene (HWG) selection

For each subregion, top-weighted genes were selected based on their average SVM weights derived from either the general model or the species-specific models. As previously described, genes with larger absolute weights contribute more significantly to the classification decision. Moreover, prior analysis indicated that approximately 600 top-weighted genes are sufficient for accurate classification of the dorsoventral axis (Figure 2I and J). Based on this, we first assigned a rank to each gene according to its weight: a gene with rank = 1 (or = -1) corresponds to the highest positive (or negative) weight, indicating the strongest contribution to the dorsal (or ventral) classification. We then defined HWGs as the top 300 genes with the highest positive (dorsal) or negative (ventral) ranks in each subregion.

## Common HWG and HDG calculation

We calculated common HWGs (Figure 4B) and HDGs (Figure 5I, J, and K) in n species. For n species in a subregion, the number of common HWGs ng is defined as the size of the union of the intersection sets of all combinations of n species selected from the five species:

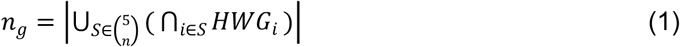

where *S* contains species from the combination of 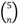, *HWG_i_*is the HWGs in species i, and |.| indicates the cardinality (i.e., the number of genes). The common HDG number was calculated similarly.

## Cross-predictions and the calculation of AUC

We performed cross-predictions of dorsoventral classifications for each species using the species-specific models from all other species in each subregion (Figure 4C and D). Since the intercept of each model does not affect the genetic network represented by the weights but does influence the accuracy when using another species’ model, we calculated the receiver operating characteristic (ROC) curves and the area under the curve (AUC) by adjusting the intercept value. This approach involves moving the hyperplane along its plane normal in the high-dimensional space. For a given intercept, the hyperplane classifies dorsal and ventral labels based on the side where each spot is located. This classification derives a true positive rate (TPR) and a false positive rate (FPR). AUC is then calculated from all TPRs and FPRs.

We also cross-predicted dorsoventral classifications for adult human snRNA-seq data from^26^ using the general models (Figure S5A and B). In this dataset, the authors annotated DG neurons and pyramidal neurons. DG neurons were predicted using the GrDG and PoDG models, while pyramidal neurons were predicted using the CA1, CA2, and CA3 models. For each model, 10 replicates of predictions were conducted. In each replicate, 1000 dorsal and 1000 ventral cells were randomly sampled for predictions. AUC was calculated in the same manner as described above.

## Numeric simulations of the ancestral confinement theory (ACT)

We conducted simulations to validate the ACT (Figure 4F, G, H, I, J, and L). The top-weighted n genes from mouse CA1 were selected, and their average scaled expressions across dorsal spots ***g_0,d_*** and ventral spots ***g_0,v_*** were calculated as the initial parameters for the ancestor. Next, we applied Brownian motion to simulate genetic drift^68,69^ for the selected genes. The probability distribution of the expression ***g_t_*** of a gene after genetic drift over time t is given by a normal distribution:

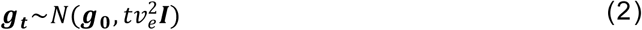

where ***g_0_*** is the *n* × 1 initial vector (i.e., the gene expression of the ancestor), ve is the evolutionary speed (kept constant for all genes), and ***I*** is the identity matrix. At a given *t* and *v_e_*, we randomly sample one value from the distribution as the evolved gene expression for each gene. This process is performed separately for dorsal and ventral spots, setting ***g_0_*** = ***g_0,d_*** and ***g_0_*** = ***g_0,v_***, resulting in the expressions ***g_t,d_*** and ***g_t,v_***, respectively. Next, we applied ancestral confinement to the evolved expressions. We projected ***g_t,d_*** onto the dorsal-side hyperplane, which is parallel to the hyperplane characterizing dorsoventral differences in the ancestor and passes through the dorsal ancestral expression. Similarly,***g_t,v_*** was projected onto the ventral-side hyperplane. This resulted in the confined expression vectors ***g_t,d,conf_*** and ***g_t,v,conf_***. Using these confined expressions, we generated *m* dorsal and *m* ventral spots from a normal distribution:

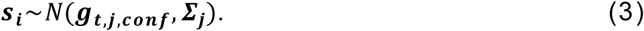

Here, ***s_i_*** is the *n* × 1 expression vector of spot *i* from *m* spots, *j ∈ {d, v}* indicates dorsoventral origin, ∑_j_ = 0.01 · *diag*(|***g_t,d,conf_***|) is the covariance matrix where |***g_t,d,conf_***| is the vector containing the element-wise absolute values of ***g_t,d,conf_*** . This generated the expression matrix for a simulated species after evolutionary time *t* . We then used SVM to determine the hyperplane for this simulated species and retrieved the weights of the n genes.

In this study, we set the number of genes *n* = 100, rationalized by the observation that a relatively small fraction of high-weighted genes is sufficient to achieve high SVM accuracy (Figure 2I and G). In this case, we defined top-weighted 10 genes for dorsal or ventral as HWGs. The number of dorsal and ventral spots *m* was set to 100 each. We simulated 10 species, each evolving at a different evolutionary speed. The simulated data of three species are visualized via UMAP. For each evolutionary time point t, we selected HWGs for each species and calculated the number of common HWGs across species. We also recorded the weights of genes at each time point. Additionally, we calculated the angles between the normal vectors of the SVM hyperplanes for each species and the ancestor. We also computed the angle between the hyperplane of the general model (trained using data from all 10 species) and that of the ancestor. This process was repeated ten times for each time point to ensure robustness.

## Dorsoventral analysis on other amniotes

In this study, we also evaluated whether dorsoventral molecular differences are conserved across a broader range of species, including rhesus macaque, chicken, lizard, and turtle (Figure 4M, N, O, P, R, and Figure S7B). Analyses were restricted to glutamatergic neurons, which represent the predominant neuronal population and display clear subregion-specific distributions.

We first defined neuron identities in each species. For the rhesus macaque, cell types were annotated using the human reference dataset from^80^ via Seurat’s label transfer pipeline^98^. Anchors were identified using the FindTransferAnchors function with parameters: normalization.method = “SCT”, recompute.residuals = FALSE, and reference.reduction = “pca”. Cells with transfer scores below 0.25 were excluded. Low-abundance cell types (< 40 cells) were manually inspected and removed. Glutamatergic neurons corresponding to CA1, CA2, CA3, and EC were extracted for downstream analysis.

For chicken, cell types were assigned based on transcriptomic similarity from^72^. Specifically, “Ex_CACNA1H_KIT” was annotated as CA1, “Ex_CACNA1H_CPA6” as CA3, and “Ex_CACNA1H_PROX1” as DG. Multiple clusters—”Ex_DACH2_LUZP2”, “Ex_DACH2_GRIK4”, “Ex_SATB2_OVOA”, “Ex_DACH2_LHX2”, “Ex_DACH2_MGAT4C”, “Ex_DACH2_NR4A3”, “Ex_DACH2_ADAMTS5”, and “Ex_SATB2_SOX6”—were annotated as EC.

For lizard, we followed the classification from^73^, where clusters “e06” and “e07” were annotated as CA1, “e09” and “e10” as CA3, and “e01”, “e02”, and “e03” as DG. Similarly, for turtle, “e35”, “e36”, “e37”, and “e38” were annotated as CA1, “e33” and “e34” as CA3, and “e29”, “e30”, “e31”, and “e32” as DG.

As a negative control, we used mouse leukocytes from^74^. As a positive control, we included mouse hippocampal and entorhinal glutamatergic neurons (CA1, CA2, CA3, DG, and EC) from^75^.

The expression data of all extracted cells were first normalized using SCTransform. Gene names were then converted to their human one-to-one orthologs; genes without a mapped ortholog were assigned an expression value of zero. Gene expressions were then projected onto the normal vectors of the SVM hyperplanes derived from the general models for each subregion. For CA1, CA2, and CA3 neurons, SVM models trained on CA1, CA2, CA3, LMol, Rad, and Or were used. For DG neurons, models from Mol, GrDG, and PoDG were used. For EC neurons, models from MECsu, MECint, and MECde were applied. For the negative control cells, all SVM models were used for projection.

As a background distribution, all cells were also projected onto 10000 randomly generated vectors. Median absolute deviations (MADs) of the projection values were calculated. One-sided permutation tests were then conducted, in which the p-value was defined as the fraction of MADs from the background distribution that exceeded the MAD from the SVM projection.

## Construction of phylogenetic trees

We constructed phylogenetic trees for each subregion across five species, with the general SVM model included as the ancestral reference (Figure 5B, C, D, E, F, and G).

Pairwise distances between species were calculated based on the SVM gene weights, using the metric 1 − ***corr***(*weigℎts*), where ***corr*** denotes the Pearson correlation coefficient between species-specific weight vectors. Hierarchical clustering was performed using the BIONJ method implemented in the R package ape^102^, based on the resulting distance matrices. The general model was designated as the root of each tree. Node support was assessed using a bootstrap approach with 1000 replicates per tree.

## Normalized rank distance (NRD)

To quantify the weight changes of reversed HDGs, we defined the normalized rank distance (NRD) (Figure 5L). For a given gene in a specific subregion and species, NRD is calculated as:

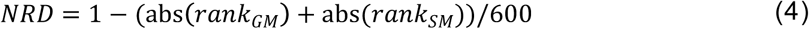

where *rank_gm_* and *rank_sm_* denote the rank of the gene in the general model and tthe species-specific model, respectively. A higher NRD indicates a stronger ranking discrepancy between the two models, emphasizing genes with substantial weight shifts.

## Consensual cell types across species

We defined the common cell types across species (Figure 6A, B, and C). The process involved the following steps. First, a k nearest-neighbor (KNN) graph was constructed using Seurat FindNeighbors with scaled data from the downsampled, integrated single-cell (nucleus) data. Louvain clustering was performed using Seurat FindClusters, with the resolution set to 0.13. Cell types were then annotated for each cluster based on known markers, including OLIGO1, OLIGO2, and MBP for oligodendrocyte (Oligo), PDGFRA and CSPG4 for oligodendrocyte precursor cells (OPC), GFAP, SLC1A2, and SLC1A3 for astrocyte (Astro), MERTK, TMEM119, and CX3CR1 for immune cells (Immune), COL1A1 and COL1A2 for fibroblast (Vascular Fibro), ACTA2 and CALD1 for smooth muscle cells (Vascular smooth), TEK and PECAM1 for endothelial cells (Vascular Endo), SLC17A6, SLC17A7, and CHN1 for excitatory neurons (Glut), SLC32A1, SLC6A1, and GAD1 for inhibitory neurons (GABA). Clusters with similar markers were merged.

Glut and GABA neurons were further subset, reintegrated, and reclustered separately to achieve finer annotations. Integration parameters were the same as those used for the whole dataset. Clustering resolution was set to 0.7 for excitatory neurons and 0.62 for inhibitory neurons. Detailed types of Glut and GABA neurons were annotated based on markers defined in mouse^75^ and human^80^. Clusters with a miscellaneous composition were labeled as “Mix”. This process resulted in 58 cell types. We then define 37 metaclasses by merging similar types of cells based on the hierarchical clustering results.

58 cell types were then compared with annotated labels from *Yao et al.* ^75^ and *Siletti et al.* ^80^ (Figure S8). Pearson correlation coefficients were calculated between the averaged PCA of our cell types and the cell types from these studies.

## Deconvolution of the spatial transcriptomes with single-cell reference

We obtained cellular compositions from spatial transcriptomic data (Figure 6D). The process involved the following steps. We first construct single-cell references for each species with the 58 cell types. To increase the representation of rare cell types, we transferred the labels from the downsampled integrated data back to all cells using the Seurat label-transfer pipeline^98^. FindTransferAnchors was used with the following parameters: normalization.method set to “SCT”, recompute.residuals set to “F”, and reference.reduction set to “pca”. Cells with transfer scores below 0.25 were excluded. We manually inspected and cleaned cell types with low cell numbers. Cell types with excessive cell numbers were downsampled to reduce computational overhead.

The Cell2location pipeline (v0. 1. 3) ^81^ was used for deconvolution. Read count matrices from single-cell (nucleus) and spatial transcriptomes, along with cell type information, were imported into Scanpy (v1.9.8)^103^ in Python (v3.9). Single-cell references for each species were constructed with default parameters. Deconvolution was performed with the following settings: N_cells_per_location set to 5, and detection_alpha set to 20.

## Analysis of cell type differences

Cell type compositions obtained from the deconvolution results were extracted and normalized such that the proportions summed to 1 within each spot. For pig and canine samples, cell types corresponding to cortical layers (Glut UL and Glut DL) were excluded prior to analysis, as these cell types are unlikely to be present in HPC. The remaining 58 cell-type clusters were aggregated into 37 metaclasses or grouped into three major categories: glutamatergic, GABAergic, and non-neural cells, for downstream analysis. To normalize and scale the compositional data, the centered log-ratio (CLR) transformation was applied to the cell type fractions, as recommended in^104^.

To compare cell type fractions along the dorsoventral axis within a given region (HPC, MEC, or subregions thereof), the average fraction for each cell type was calculated by averaging across all dorsal or ventral spots in that region. The log fold change of a given cell type was defined as the difference in average CLR values between dorsal and ventral spots. Statistical comparisons were performed using CLR-transformed fractions between dorsal and ventral spots (Figure 6 E and F). Due to the large sample size, the adjusted p-values remained extremely small even after Benjamini-Hochberg (BH) correction. To reduce the false positive rate, we further applied a more stringent threshold for significance based on differences in CLR values. Specifically, a result was considered significant only if the BH-adjusted p-value was smaller than a threshold p_0_, which was defined as the largest adjusted p-value among all comparisons where the absolute CLR difference exceeded 5.5 times the median of all absolute CLR differences.

## Acknowledgments

We thank Zexian Zeng’s lab, Ge Gao’s lab, and Zhiyuan Li’s lab for the help in data processing and analysis. This work was supported by the National Key R&D Program of China (2019YFA0802400), the Lingang Laboratory, Grant No. LG-TKN-202204-01, the State Key Laboratory of Membrane Biology (open project for Chenglin Miao), and the Qidong-SLS Innovation Fund (Grant No. 2023002029 and 2020001540). We thank the Core Facilities of the School of Life Sciences, the State Key Laboratory of Membrane Biology at Peking University for assistance with sample preparation.

## Author contributions

C.M. planned and initiated the research, supervised the project, secured funding, and designed the experimental component. J.L. designed the theoretical data-analysis component. H.Y. assisted with access to human samples. M.P. contributed to research planning. S.C. contributed to study design and performed experiments, including sample collection, histology, and transcriptomic sequencing, and also assisted with data analysis. Q.W. contributed to study design and conducted data analyses, including data cleaning, statistical analysis, machine learning, theoretical model construction, and numerical simulations. Y.F. assisted with human sample preparation. C.C., S.U., and Y.L. assisted in performing experiments. Y.Y. and K.G. assisted with data analysis. S.C., Q.W., C.C., M.P., J.L., and C.M. contributed to manuscript preparation.

## Declaration of interests

The authors declare no competing interests.

## Notes

### Competing Interest Statement

The authors have declared no competing interest.

