## Supplemental Information for "Transcriptomic characteristics along the longitudinal axis of the hippocampus and medial entorhinal cortex across species"

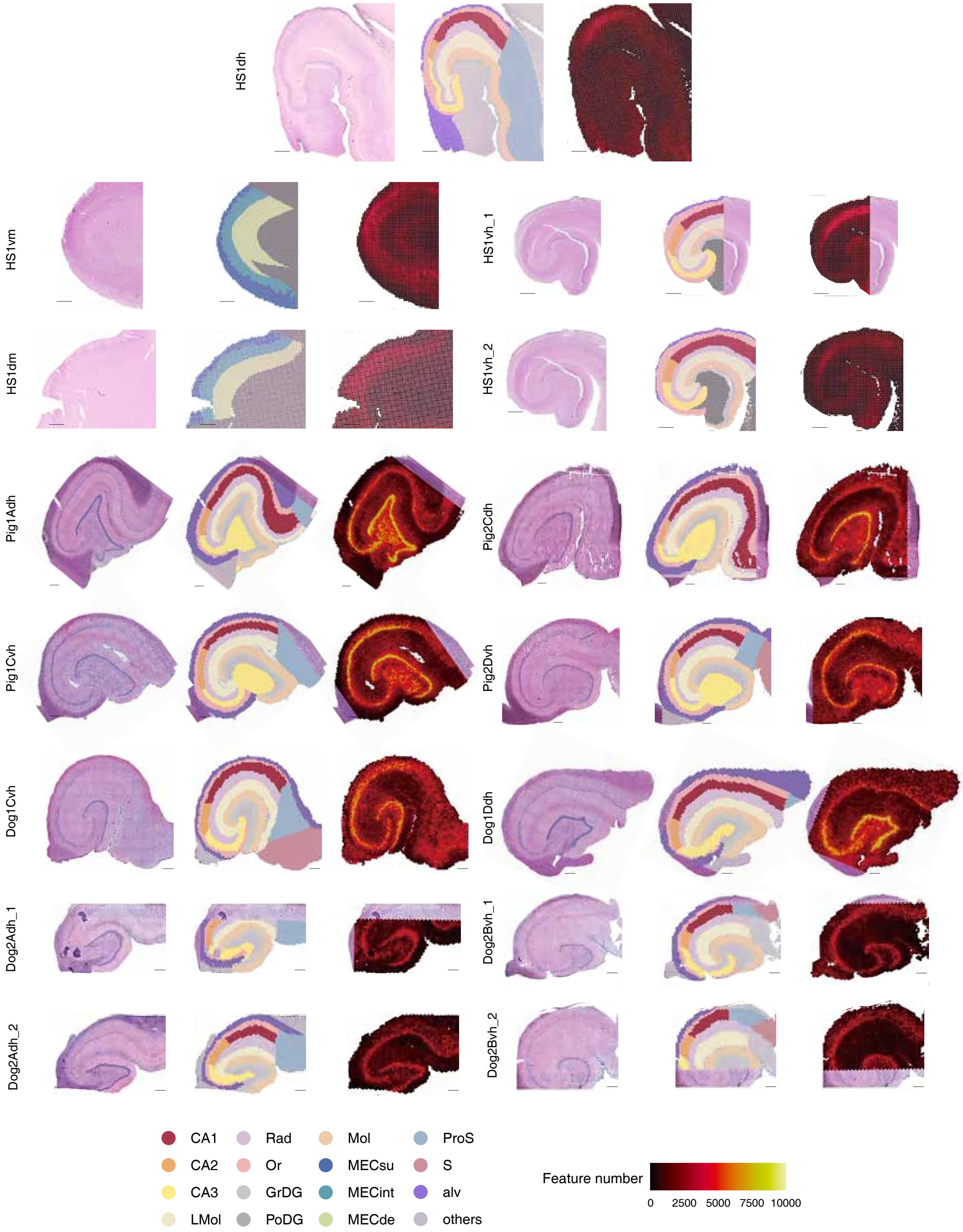


FIG. S1: **Samples slides of human, pig, and canine.** Each sample contains figures of the H.E. staining image (left), spots labeled with subregions (middle), and spots labeled with detected genes number (right). Black bar represents 500 $\mu$m.

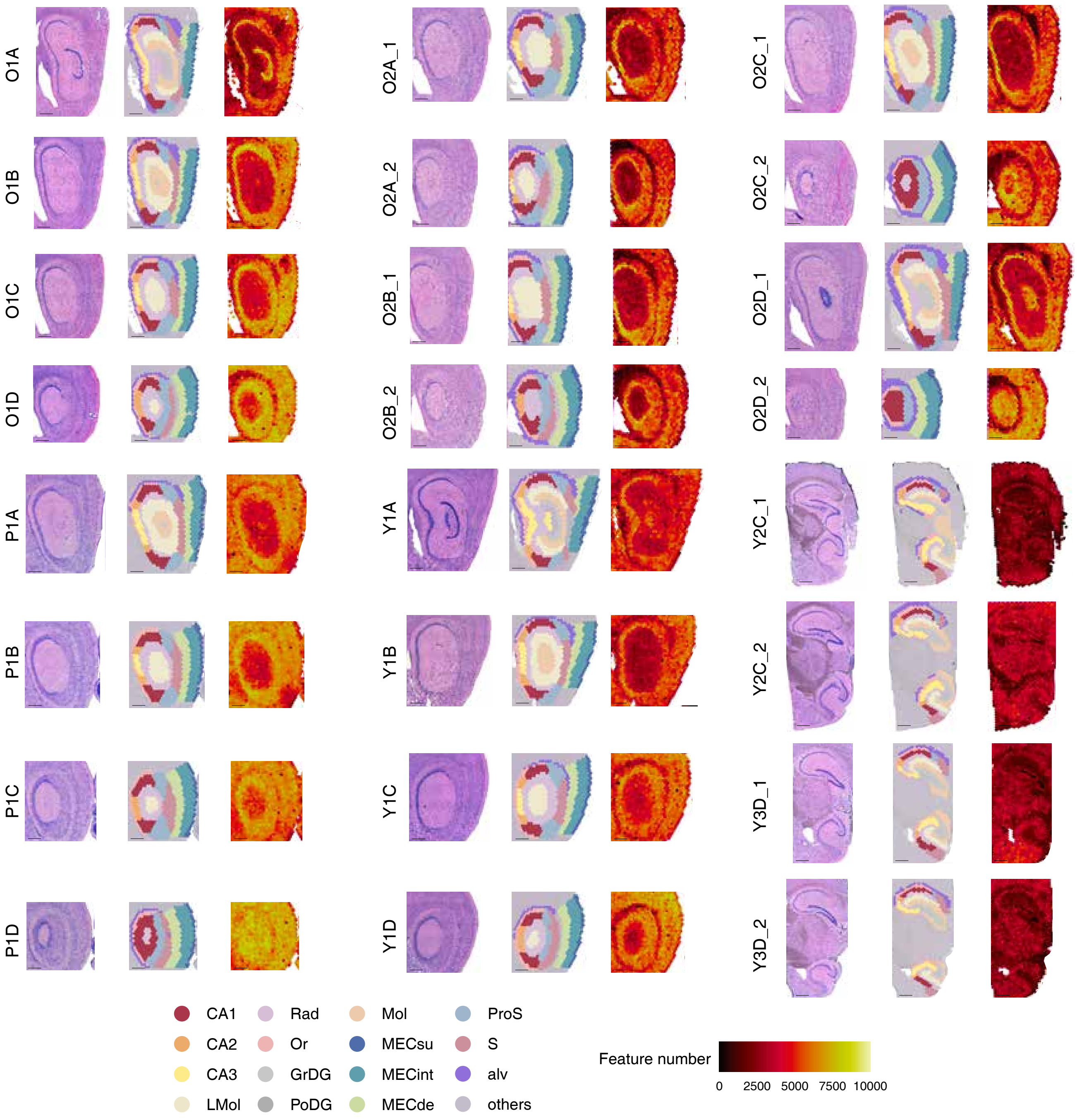


FIG. S2: **Samples slides of mouse.** Each sample contains figures of the H.E. staining image (left), spots labeled with subregions (middle), and spots labeled with detected genes number (right). Black bar represents 500 $\mu$m.

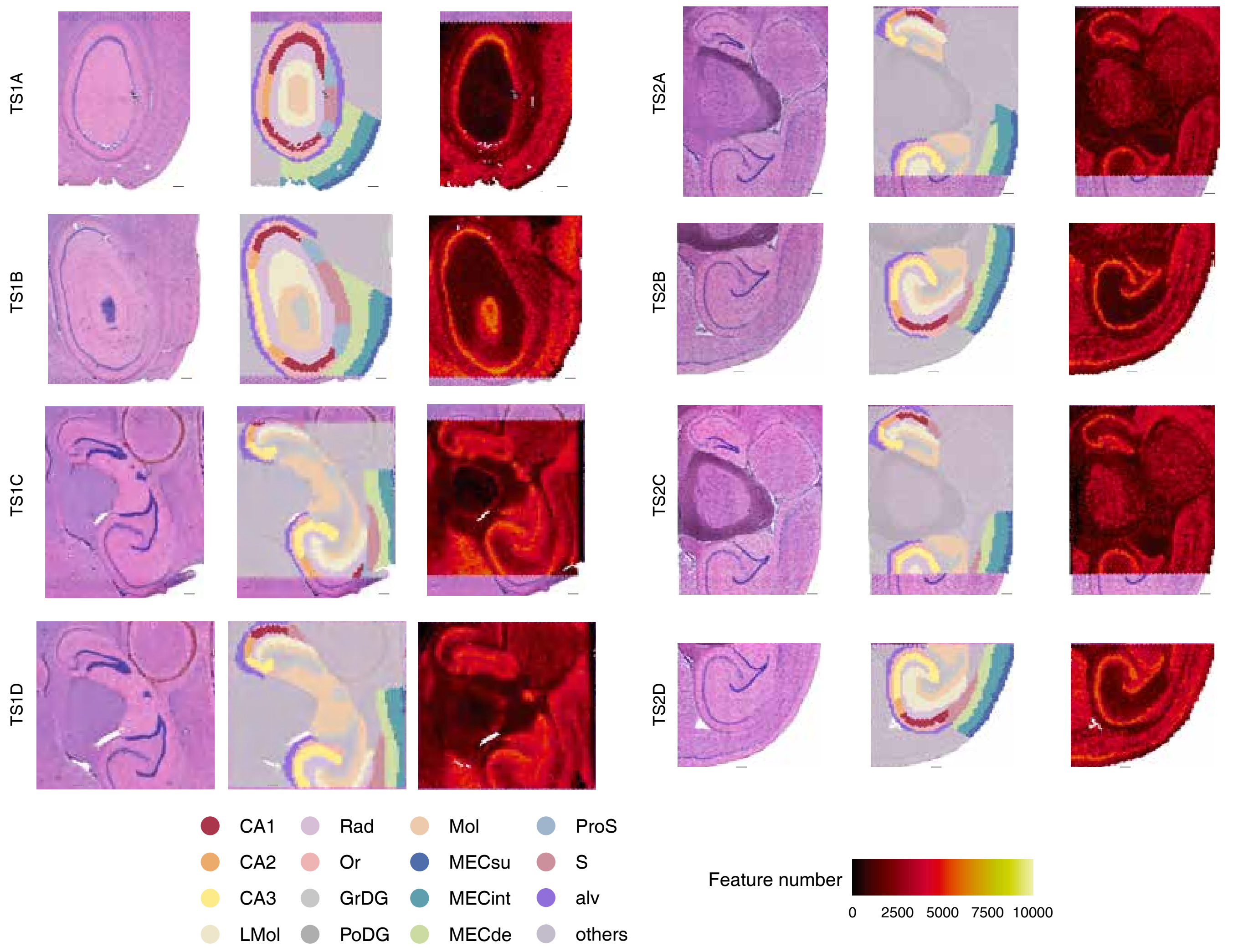


FIG. S3: **Samples slides of treeshrew.** Each sample contains figures of the H.E. staining image (left), spots labeled with subregions (middle), and spots labeled with detected genes number (right). Black bar represents 500 $\mu$m.

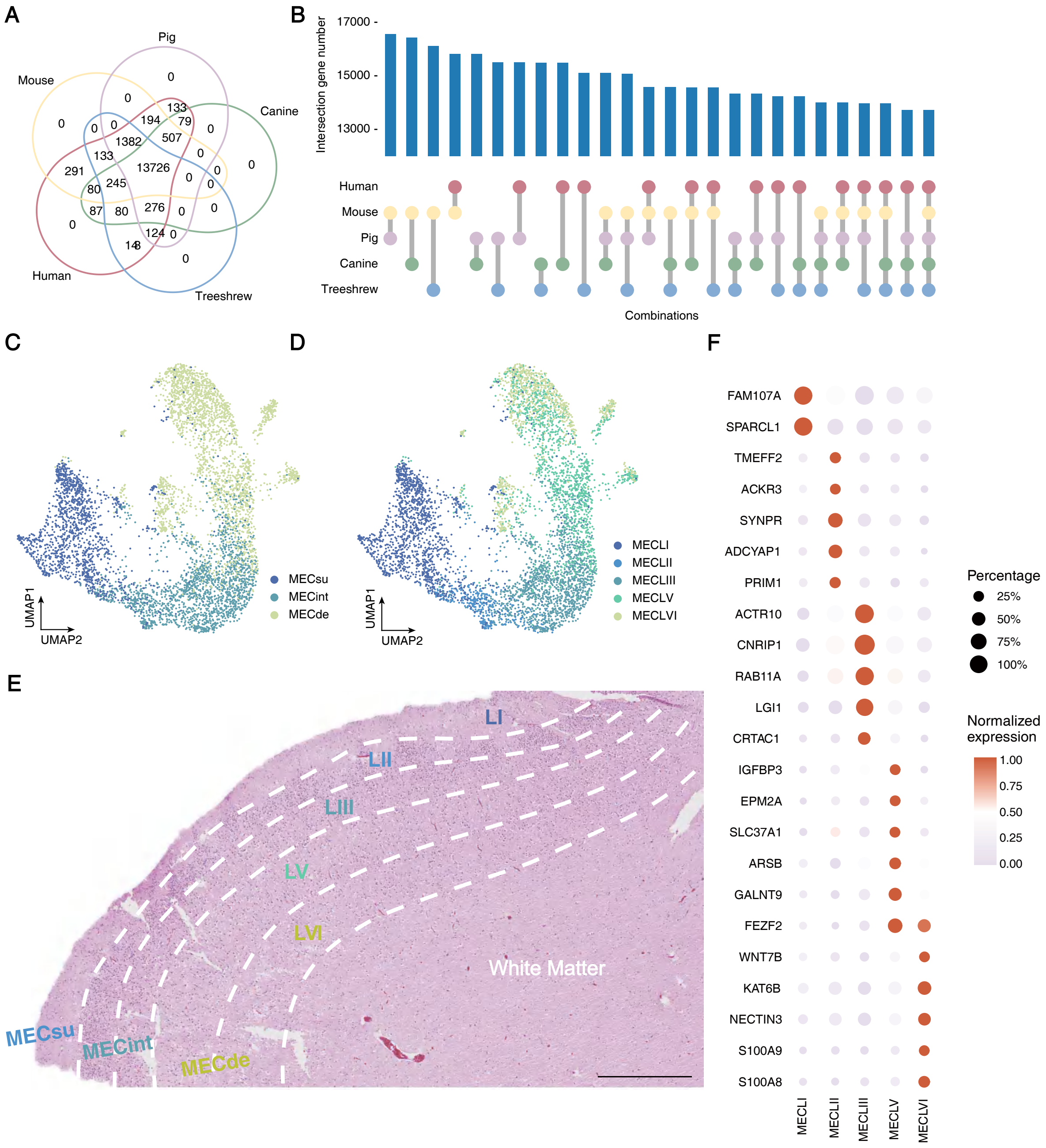


FIG. S4: **Orthologous genes and human MEC features.** Related to Figure 1 in the main text. (A) Intersections of orthologous genes among all species. (B) Intersections of orthologous genes shared across n species. (C) UMAP representation of human MEC spots colored by a 3-layer classification. (D) UMAP representation of human MEC spots colored by a 5-layer classification. (E) Anatomical representation of human MEC with both 3-layer and 5-layer classifications labels. Scale bar: 500 $\mu$m. (F) Expression of marker genes corresponding to the five layers of human MEC.

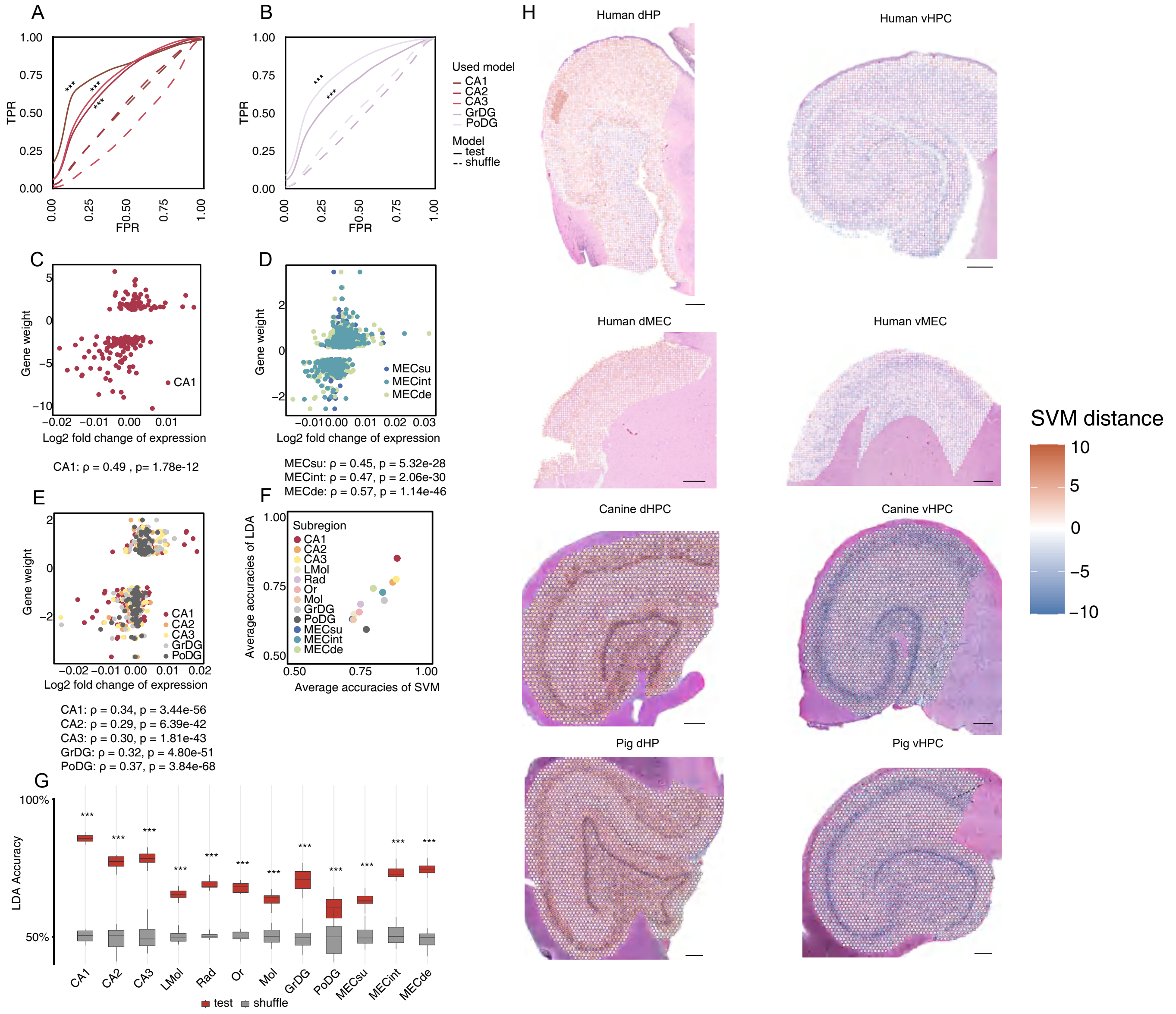


FIG. S5: **Validation of the general SVM model.** Related to Figure 2 in the main text. (A) Area Under the Curve (AUC) curves for dorsoventral classification predictions of single-nucleus RNA-seq data for human pyramidal neurons from [1] using the general SVM models from this study. Curves are smoothed across 10 replicates, where a randomly sampled set of cells is used for predictions in each replicate. Two-sided Wilcoxon tests were performed to compare AUC values between the test and shuffled groups. Significance levels are indicated with p-values as follows: "$***$"represents $p < 1e-3$. (B) Similar analysis as in (A), performed for human DG neurons from [1]. (C) Correlation between the weights of the general model for CA1 and the log2 fold changes of DEGs identified in mouse CA1 from [2]. Correlations were tested using two-sided t-tests, with the correlation coefficient and p-value shown on the right. (D) Correlation between the weights of the general models for CA1, CA2, CA3, GrDG, and PoDG and the log2 fold changes of DEGs identified in the whole mouse HPC from [3], similar to (C). (E) Correlation between the weights of the general models for MECsu, MECint, and MECde and the log2 fold changes of DEGs identified in the whole mouse MEC from [4], similar to (C). (F) Comparison of the correlation between the highest prediction accuracies of SVM and Linear Discriminant Analysis (LDA) across subregions. The Pearson correlation coefficient. Correlations were tested using two-sided t-tests ($\rho = 0.86$, p value $= 3.10e-4$). (G) LDA classification accuracies for each subregion. Two-sided Wilcoxon tests compare the accuracies between testing and shuffled groups. P-values are corrected using the false discovery rate (FDR), and significance levels are indicated by asterisks, with "***" representing FDR$<0.001$. (H) SVM distance for HPC of human, canine, and pig.

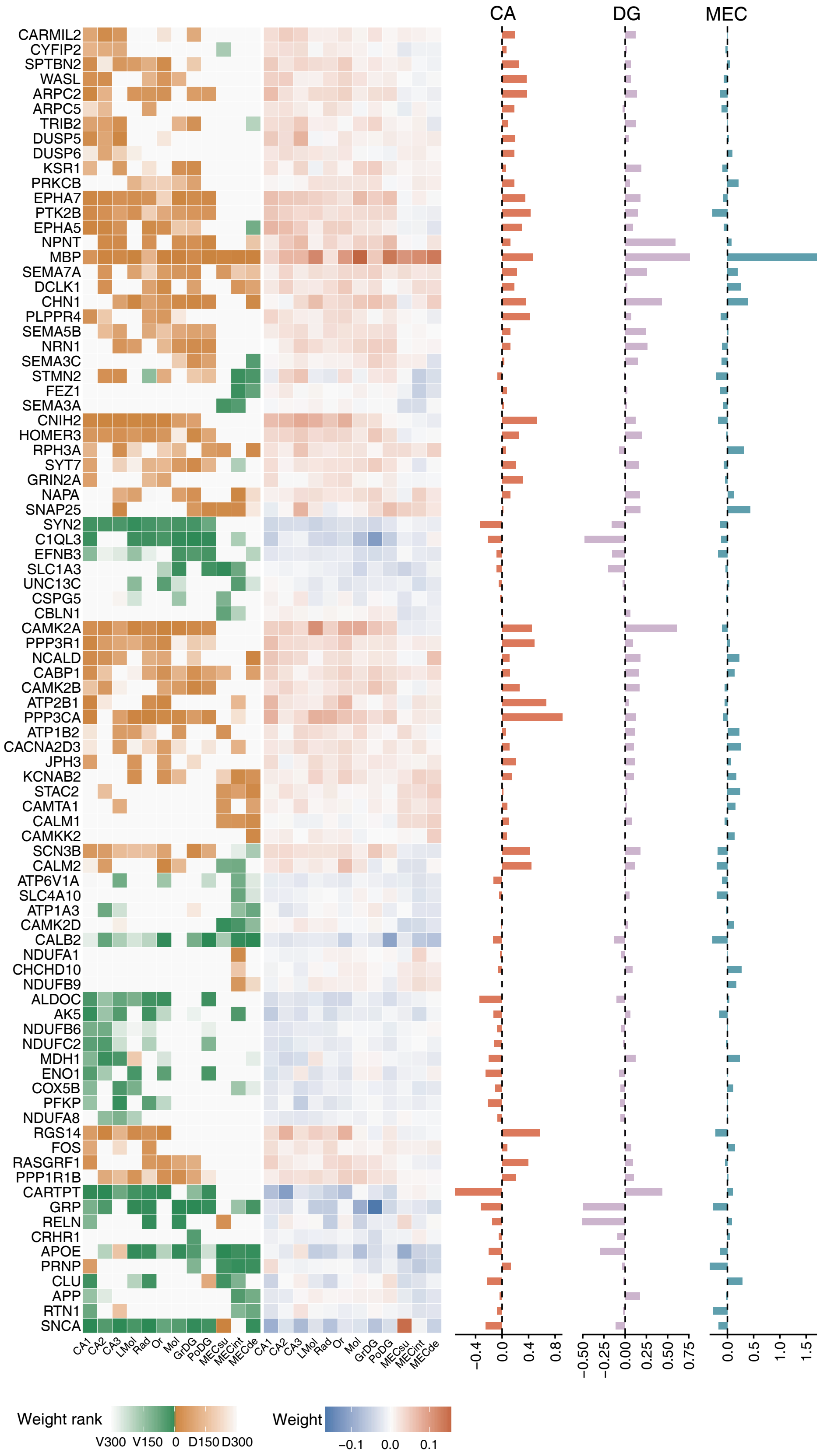


FIG. S6: **SVM weights and dorsoventral expression differences of HWGs.** Related to Figure 3 in the main text. Left: the heatmap shows the ranks (left) and SVM weights (middle) of HWGs listed in Figure 3 from the main text. The accompanying bar plots (right) display the average dorsoventral expression differences of the corresponding genes across all species in the CA, DG, and MEC regions. Positive values indicate higher expression in the dorsal region, while negative values indicate higher expression in the ventral region.

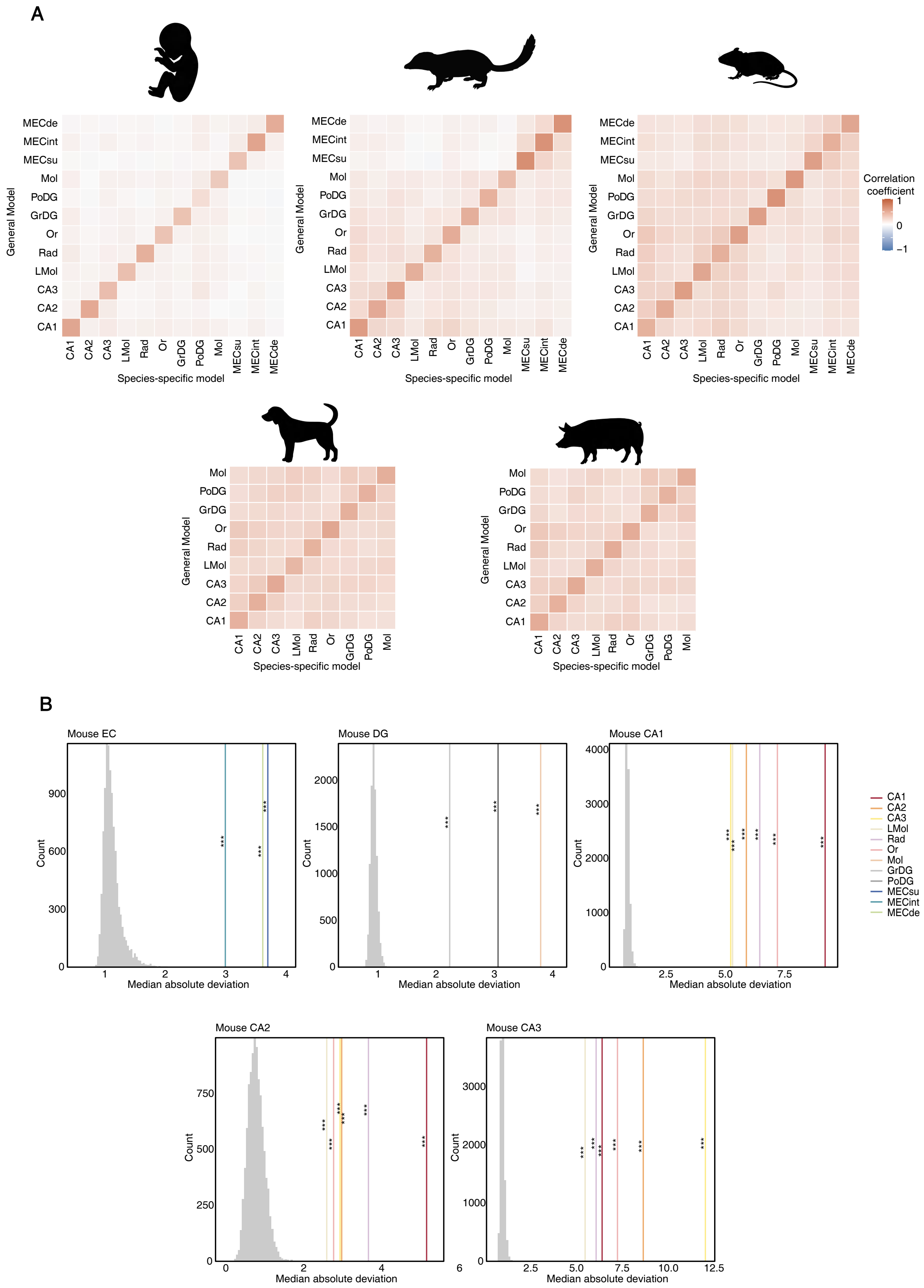


FIG. S7: **Correlation analysis and positive control for variation analysis.** Related to Figure 4 in the main text. (A) Correlation analysis between gene weights of species-specific and general SVM models. Colors represent Pearson correlation coefficients. All correlations were statistically significant with p-values < 0.05 (two-sided t-test). (B)Projection of glutamatergic neuron expression data from the mouse HPC and MEC onto the normal vectors of hyperplanes defined by the corresponding general SVM models used as positive control distributions, compared against projections onto 10000 randomly generated directions.

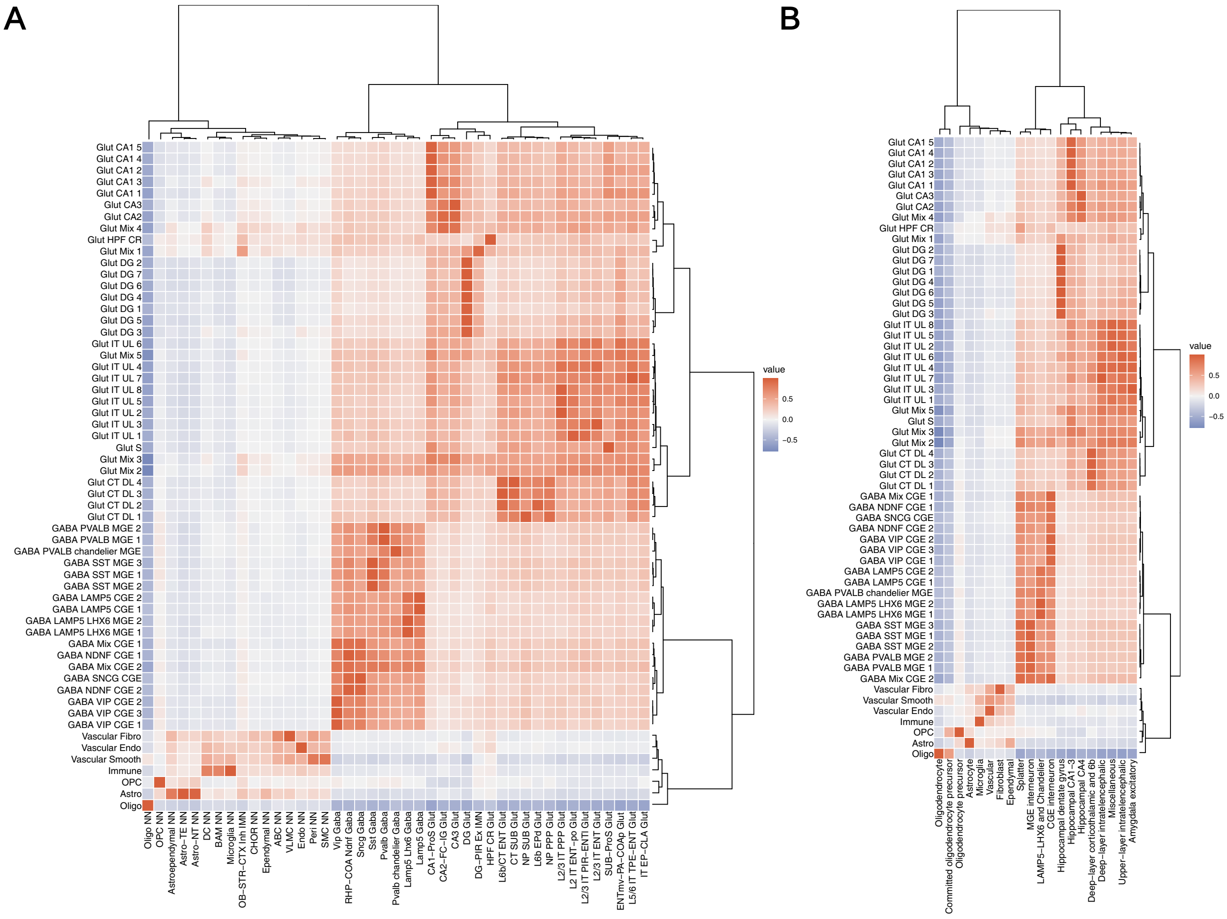

FIG. S8: **Comparisons of cell types defined in this work and in other works.** Related to Figure 4 in the main text. (a) Correlations between cell types defined in this work (y-axis) and those from [5] (x-axis). (b) Correlations between cell types defined in this work (y-axis) and those from [6] (x-axis).
